# Influenza A infection accelerates disease-associated microglia formation during physiological aging

**DOI:** 10.64898/2025.12.11.693336

**Authors:** Rogan A. Grant, Constance E. Runyan, Grace A. Minogue, Joshua S. Stoolman, Satoshi Watanabe, Fei Chen, Karolina J. Senkow, Luisa Cusick, Maxwell J. Schleck, Radmila Nafikova, Ziyan Lu, Elsie G. Bunyan, Hiam Abdala-Valencia, Jacob I. Sznajder, Robert Vassar, Rudolph Castellani, Changiz Geula, Tamar Gefen, Navdeep S. Chandel, Alexander V. Misharin, G.R. Scott Budinger

**Affiliations:** Division of Pulmonary and Critical Care Medicine, Department of Medicine, Feinberg School of Medicine, Northwestern University, Chicago, IL, USA; Mesulam Institute for Cognitive Neurology and Alzheimer’s Disease, Northwestern University Feinberg School of Medicine, Northwestern University, Chicago, IL, USA; Department of Psychiatry and Behavioral Sciences, Northwestern University Feinberg School of Medicine, Northwestern University, Chicago, IL, USA; Department of Neurology, Feinberg School of Medicine, Northwestern University, Chicago, IL, USA; Department of Pathology, Division of Neuropathology, Northwestern University Feinberg School of Medicine, Chicago, IL, USA; Simpson Querrey Lung Institute for Translational Science, Northwestern University, Chicago, IL, USA; Chan Zuckerberg Biohub, Chicago, IL, USA

## Abstract

Severe pneumonia is associated with an increased risk of cognitive decline and dementia, particularly in the elderly. Changes in microglia, the most abundant immune cell population in the brain, are also associated with cognitive decline and dementia, including the emergence of a transcriptional cell state referred to as disease-associated microglia (DAM). We sought to test the hypothesis that non-neuroinvasive influenza A virus (IAV) pneumonia results in transcriptional responses in brain microglia that drive premature expansion of DAM. Using bulk and single-cell RNA-sequencing, metabolomics, and spatial transcriptomics, we profiled neuroimmune populations in young, middle-aged, and old male mice during IAV infection and recovery. We observed an increased abundance of DAM, interferon-responsive microglia (IRM), CD4+ T cells, and CD8+ T cells in white matter regions beginning in middle age and persisting in old animals, irrespective of IAV infection. DAM exhibited a metabolic shift toward aerobic glycolysis with disrupted TCA cycling, citrulline depletion, and an elevated itaconate/α-ketoglutarate ratio. Spatial transcriptomic profiling of the human middle frontal gyrus (MFG) in normal agers, SuperAgers, and patients with dementia revealed an analogous accumulation of DAM and CD8+ T cells in white matter. IAV pneumonia induced a transient immunosenescent-like response in microglia, marked by glucocorticoid-responsive gene expression and *Ccnd3* upregulation. In response to IAV pneumonia, DAM expanded in middle-aged mice, whereas old mice were elevated at baseline and were largely unaffected by IAV infection. The age-related expansion of DAM was unaffected by pharmacological depletion and repopulation of microglia with a CSF1R antagonist or genetic gain or loss of function of the phagocytic receptor MERTK, suggesting the DAM phenotype is driven by the CNS microenvironment, rather than cell-intrinsic mechanisms. Our findings suggest that IAV pneumonia induces an acute immunosenescence response in microglia and accelerates the age-dependent expansion of DAM in white matter.

## Introduction

Pneumonia is the most common cause of death from an infectious disease worldwide and disproportionately affects the elderly. Survivors of severe pneumonia requiring hospitalization have a substantially increased risk of cognitive decline or dementia that persists for at least 3 years after hospitalization. Longitudinal analysis suggests that pneumonia accelerates cognitive decline in individuals already exhibiting mild cognitive impairment, with associations becoming significantly stronger with age^1–3^. Cognitive decline after pneumonia has become an important global public health concern as the SARS-CoV-2 virus pandemic resulted in a sharp increase in the number of severe pneumonia survivors, many of whom experience lasting effects on cognition^4–6^. The biological mechanisms responsible for cognitive decline in pneumonia survivors and their links to the biology of aging, however, are largely unknown.

As very few pneumonia pathogens exhibit tropism for the brain, investigators have suggested that endocrine signals from the infected lung, capable of crossing the blood-brain barrier, are responsible for cognitive decline after pneumonia. Pro-inflammatory cytokines are known to cross the blood-brain barrier and gain access to the CNS, where they can be sensed by CNS-resident immune cells, including microglia^1,3,7–10^. Changes in microglia and neurons have been observed during acute infection of rodents with influenza viruses and SARS CoV-2^11–15^. We and others have observed microglial activation in the brains of patients who died following SARS-CoV-2 infection in the absence of direct viral infection of the CNS^16–21^.

Microglial cell states have been associated with age-related cognitive decline and dementia in humans and mouse models. Notable among these are disease-associated microglia (DAM) and interferon-responsive microglia (IRM). These cell states accumulate during normal and successful aging and are relatively expanded in patients with dementia, including Alzheimer’s disease, and in murine models of dementia^22–27^. Some have suggested that aging may “prime” the neuroimmune system for an aberrant response to peripheral immune activation,^28–31^ and enhancement of pathological hallmarks has been observed in mouse models of neurodegeneration with peripheral inflammatory stimuli^32,33^.

We sought to test the hypothesis that influenza A virus (IAV) pneumonia accelerates the expansion of DAM in normal aging. We phenotyped neuroimmune cells over the course of IAV pneumonia and recovery in young adult, middle-aged, and old mice using bulk and single-cell RNA sequencing, metabolomics, and spatial transcriptomics on both mouse and human brain samples. We observed age-related neuroinflammation in microglia and CD8+ T cells in both mouse and human postmortem brains, enriched in white matter. Irrespective of age, we identified a transient inflammatory response in microglia, characterized by a glucocorticoid response signature and markers of cell-cycle arrest. In middle-aged mice specifically, DAM rapidly expanded during acute IAV pneumonia and persisted after recovery. Our findings suggest IAV pneumonia induces an acute immunosenescence response in microglia and affects the CNS microenvironment to accelerate the age-dependent expansion of DAM in white matter.

## Results

### Physiological aging is associated with microglia activation and T cell invasion in the brain parenchyma

We used definitions for murine age from the Jackson Laboratory, which defines mature adult / young adult mice as 3-6 months old, middle-aged mice as 10-14 months old, and old mice as 18-24 months of age^34^. We performed bulk RNA-seq on flow cytometry-purified microglia from old and young adult mice (Supp. Fig. 1A; Supp. Table S1). We observed widespread transcriptomic remodeling in microglia from old animals, characterized by upregulation of key markers of DAM, including *Spp1*, *Itgax*, *Axl*, *Cst7*, *Gpnmb*, *Clec7a*, *Lpl*, *B2m*, *App*, and *Apoe*^22,23,35^; markers of IRM, including *Ifit2*, *Ifit3*, *Ifit3b*, *Ifi204*, *Ifi44*, *Ifi27l2a*, *Oas1a*, *Oas2*, *Oasl2*, *Oas3*, *Mx1*, and *Ccl12* ^24,35^; and genes involved in microglial proliferation, including *Csf1*, and *Mki67*. We observed downregulation of genes associated with homeostatic function in microglia from old mice, including *Mertk*, *Tmem119*, *P2ry12*, *Cx3cr1*, *Slco2b1*, and *Csf1r*. Many of the differentially expressed genes suggested a change in microglial metabolism with advancing age, including upregulation of genes encoding enzymes involved in glycolysis (*Pkm*, *Pgk1*, *Pgam1*), as well as fatty acid beta oxidation and lipid homeostasis (*Acacb*, *Lpl*, *Apoe*). *Ass1*, encoding arginosuccinate synthase-1, was among the most upregulated genes with advancing age. ASS1 catalyzes the synthesis of arginosuccinate, a precursor of arginine, from citrulline and aspartate as part of the urea cycle and has been linked to inflammatory activation in macrophages (Fig. 1a; Extended Data 1)^36^. Gene-set enrichment analysis (GSEA), using DAM signature genes,^22^ Sennet-Mayo senescence gene list (Senmayo)^37,38^, and hallmark gene sets from MSigDB^39^, further highlighted these changes, with significant enrichment of DAM marker genes, the interferon alpha response and interferon gamma response, cholesterol homeostasis, glycolysis, hypoxia, mitotic spindle assembly, and complement pathway activation. Enrichment of NF-κB-responsive gene expression, G2M checkpoint, and Senmayo was also suggestive of an immunosenescent-like phenotype (Fig. 1b).

**Figure 1.**
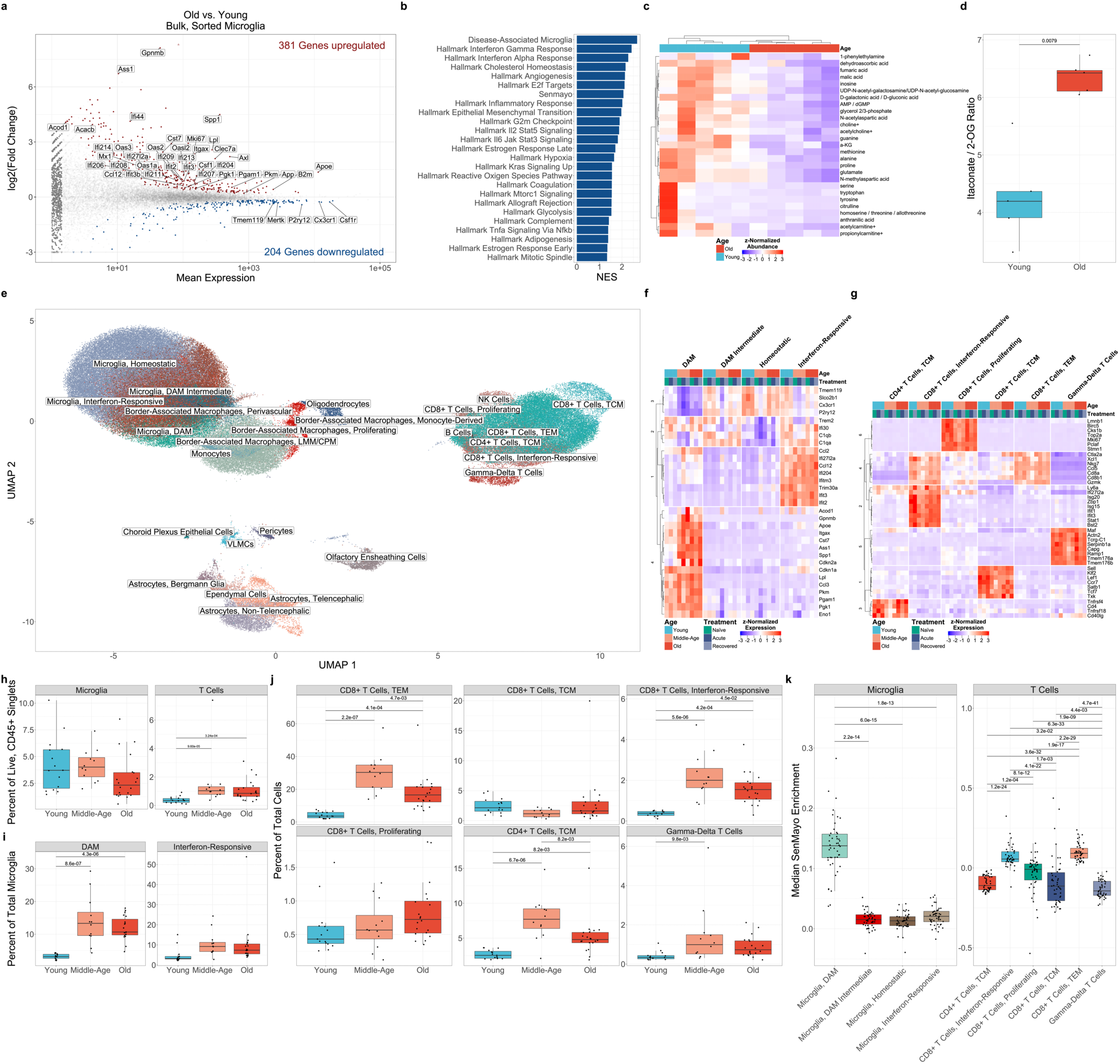
Physiological aging is associated with microglia activation and lymphocyte invasion in the brain parenchyma. **(a)** MA plot of differentially expressed genes in flow-cytometry sorted microglia from old (20-21mo; n = 9) vs young adult mice (5-7mo; n = 8). Significantly (q < 0.05, Wald test) upregulated genes are shown in red, and significantly downregulated genes are shown in blue. Genes shown in gray are not significantly differentially expressed. Genes represented by triangles are outside the plot limits. **(b)** Gene-set enrichment analysis (GSEA) of significantly enriched gene lists (q < 0.05, permutation test on Monte-Carlo simulations) from bulk RNA-seq in (a) **(c)** Hierarchical clustering of z-normalized metabolite abundance of analytes from flow-cytometry sorted microglia from old (18mo; n = 5) and young adult (4mo; n = 5) mice with nominally significant differences (p < 0.05; t-tests). **(d)** Peak area ratios of itaconate / 2-oxoglutarate (2-OG) in bulk microglia metabolomics from (c). Comparison was made with the Wilcoxon rank sum test. **(e)** UMAP of scRNA-seq results from 167,598 flow-cytometry sorted neuroimmune cells passing quality filters isolated from whole mouse brains of C57BL/6JN mice in the steady state (naïve), during acute IAV infection (1 week post-infection), and after recovery from pneumonia (5 weeks post-infection) in young adult (4-5 months old; n_naïve_ = 5, n_acute_ = 5, n_recovered_ = 5), middle-aged (12-13 months old; n_naïve_ = 3, n_acute_ = 4, n_recovered_ = 5), and old (18-19 months old; n_naïve_ = 4, n_acute_ = 8, n_recovered_ = 8) **(f-g)** Hierarchical clustering of mean marker gene expression in microglia **(f)** and T cells **(g)** by cell state, age and treatment from (e). **(h)** Relative flow cytometric abundances of microglia and T cells from single-cell suspensions as a proportion of live, singlet, CD45^+^ cells from whole mouse brains of C57BL/6JN mice in the steady state (naïve), during acute IAV infection (1 weeks post-infection), and after recovery from pneumonia (5 weeks post-infection) in young adult (4-5 months old; n_naïve_ = 5, n_acute_ = 5, n_recovered_ = 5), middle-aged (12-13 months old; n_naïve_ = 3, n_acute_ = 4, n_recovered_ = 5), and old age (18-19 months old; n_naïve_ = 4, n_acute_ = 8, n_recovered_ = 8). Comparisons are Wilcoxon rank sum tests with FDR correction. **(i-j)** Relative abundances of select microglia **(i)** and T cell **(j)** states as a percentage of total microglia in scRNA-seq dataset from (e). Comparisons are t-tests with FDR correction. **(k)** Median SenMayo module enrichment by mouse and cell state in microglia and T cells from scRNA-seq dataset in (e).

To further explore the metabolic alterations observed in microglia at the transcript level, we performed unbiased profiling of polar metabolites in flow cytometry-purified microglia from young adult and old mice (Supp. Fig. 1b; Supp. Table S2). As suggested by our transcriptomics analysis, we observed a marked reduction in many TCA cycle metabolites in old animals, including fumarate, malate, and α-ketoglutarate; as well as TCA products, with particularly strong depletion of amino acids and their derivatives. We also observed age-related decreases in metabolites involved in purine synthesis and choline metabolism. Notably, citrulline, the substrate for ASS1, was depleted in microglia from old animals, consistent with functional upregulation of ASS1 (Fig. 1c)^36^. We also observed a significant increase in the itaconate / α-ketoglutarate ratio in microglia from old animals, consistent with activation of inflammatory programs in macrophages that require TCA cycle disruption (Fig. 1d)^40–43^. This finding was further supported by a strong upregulation of *Acod1* (*Irg1*), encoding the enzyme responsible for itaconate synthesis.^44^

We measured the impact of ongoing age-related neuroinflammation on the neuroimmune response to IAV pneumonia. Old, compared to young adult or middle-aged mice, exhibit increased mortality and delayed recovery from IAV pneumonia, providing a model that recapitulates the age-related susceptibility to viral pneumonia observed in elderly^45^. Similar to humans, sialic acid receptors necessary for IAV entry are only expressed in the airway and alveolar epithelium of mice.^46^ Hence, intratracheal infection with IAV does not result in viremia or extrapulmonary infection, including in the brain^47^, thereby excluding a direct impact of viral infection on any resulting changes in the CNS. Accordingly, we intratracheally infected cohorts of young adult, middle-aged, and old mice with an inoculum of IAV, titrated to induce similar weight loss and mortality in each age group (average weight loss 21±9% of starting weight and average mortality 42%; Supp. Fig. 1c-d)^45,48^. We harvested the brains from each cohort in naïve animals, during acute infection (1 week post-infection), and after recovery (5 weeks post-infection) and flow-cytometry sorted brain macrophages, T cells, and astrocytes and analyzed them using scRNA-seq (Supp. Fig. 1e; Supp. Table S3).

We resolved homeostatic microglia, DAM^22,23,35^, IRM^24,35^, border-associated macrophages (BAM), lymphocytes, astrocytes, oligodendrocytes, and structural cells (Fig. 1e; Extended Data 2). We also identified an intermediate DAM-like state characterized by elevated *Trem2* expression^22^. Both IRM and intermediate DAM expressed genes encoding proteins in the complement pathway, including *C1qa* and *C1qb*. DAM were the primary contributor to the SenMayo signature observed in bulk RNA-seq, with elevated expression of both *Cdkn1a* (encoding p21) and *Cdkn2a* (encoding p16). Notably, DAM appeared to be the primary contributors to the metabolic shifts observed in both bulk RNA-seq and metabolomics on these cells, exhibiting increased expression of *Pkm*, *Pgk1*, *Pgam1*, *Lpl*, *Apoe*, *Ass1*, and *Acod1* (Fig. 1f).

Within T cells, we identified effector memory CD8+ T cells (TEM), which expressed genes associated with activation and local expansion (*Gzmk*, *Ccl5*, and *Nkg7*). In addition, we identified a cluster of CD8+ T cells that express interferon-responsive genes, including *Ifit1*, *Ifit3*, *Stat1*, and *Isg20*, which we annotated “Interferon-Responsive”. Finally, we identified a cluster of proliferating CD8+ T cells expressing markers of cell division, including *Mki67* and *Top2a* (Fig 1g).

Using flow cytometry, we detected an increased abundance of total T cells in the brains of both middle-aged and old animals, compared to young adult animals. Total numbers of microglia did not change with advancing age (Fig. 1h; Supp. Fig. 1e). Using scRNA-seq abundance data, we identified increased abundance of DAM and IRM, CD8+ T effector memory (TEM), interferon-responsive CD8+ T cells, CD4+ T central memory (TCM) and γδ T cells in middle-age, which remained elevated in old animals. An increased abundance of proliferating CD8+ T cells was observed only in old animals (Fig 1i,j). Notably, many of these age-associated cell states, including DAM, CD8+ TEM, and interferon-responsive CD8+ T cells were significantly enriched for genes included in Sen-Mayo score (Fig. 1k). The aging mouse brain is therefore characterized by accumulating sterile inflammation originating from both resident microglia and invading T lymphocytes.

### Age-related neuroinflammation is localized to white matter

Our findings suggest normal aging is associated with an expansion of DAM and IRM and T cell invasion into the brain parenchyma that is detectable by middle age. To better understand the spatial localization of these cells in the aging brain, we performed single-cell spatial transcriptomics on whole, decalcified mouse skulls with intact meninges. We resolved subtypes of neurons, glia, immune cells, and structural cells to provide an atlas of the aging mouse brain and meninges at subcellular resolution (Fig. 2a; Extended Data 3). We detected heterogeneous populations of microglia and T cells, including DAM, IRM, and CD8+ T cells (Supp. Fig. 2a, c-e). Similar to flow cytometry and single-cell RNA-seq, age-related increases were again observed in all three cell states (Fig. 2b-c). Consistent with previous reports, DAM were strongly enriched in white matter regions (Fig. 2d; Supp. Fig. 2b)^25,26,49^. IRM and CD8+ T cells were diffusely distributed throughout the brain parenchyma with enrichment in white matter regions. We identified CD8+ T cells in the brainstem, hypothalamus, olfactory regions, and hippocampus in old animals (Fig. 2e).

**Figure 2.**
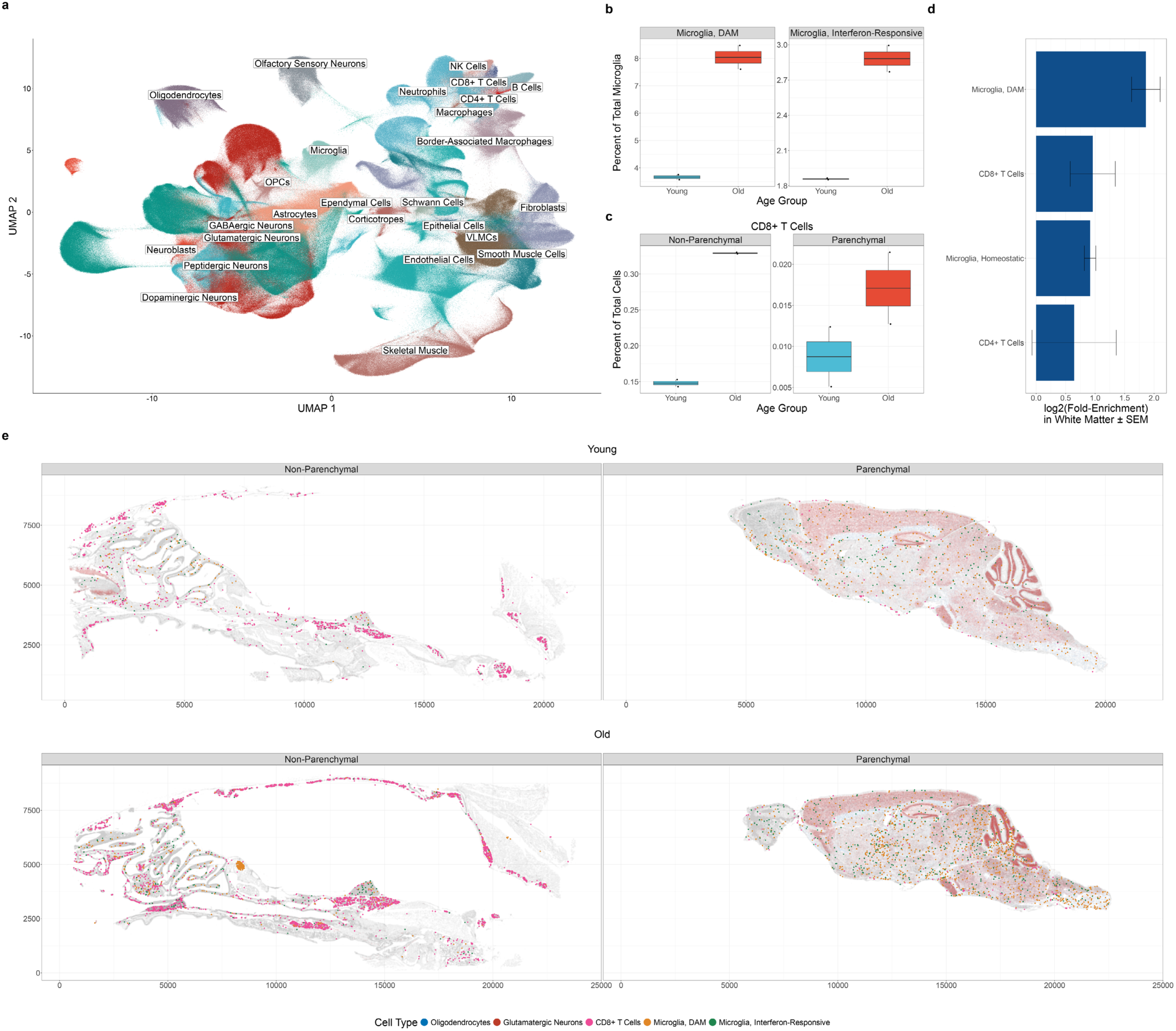
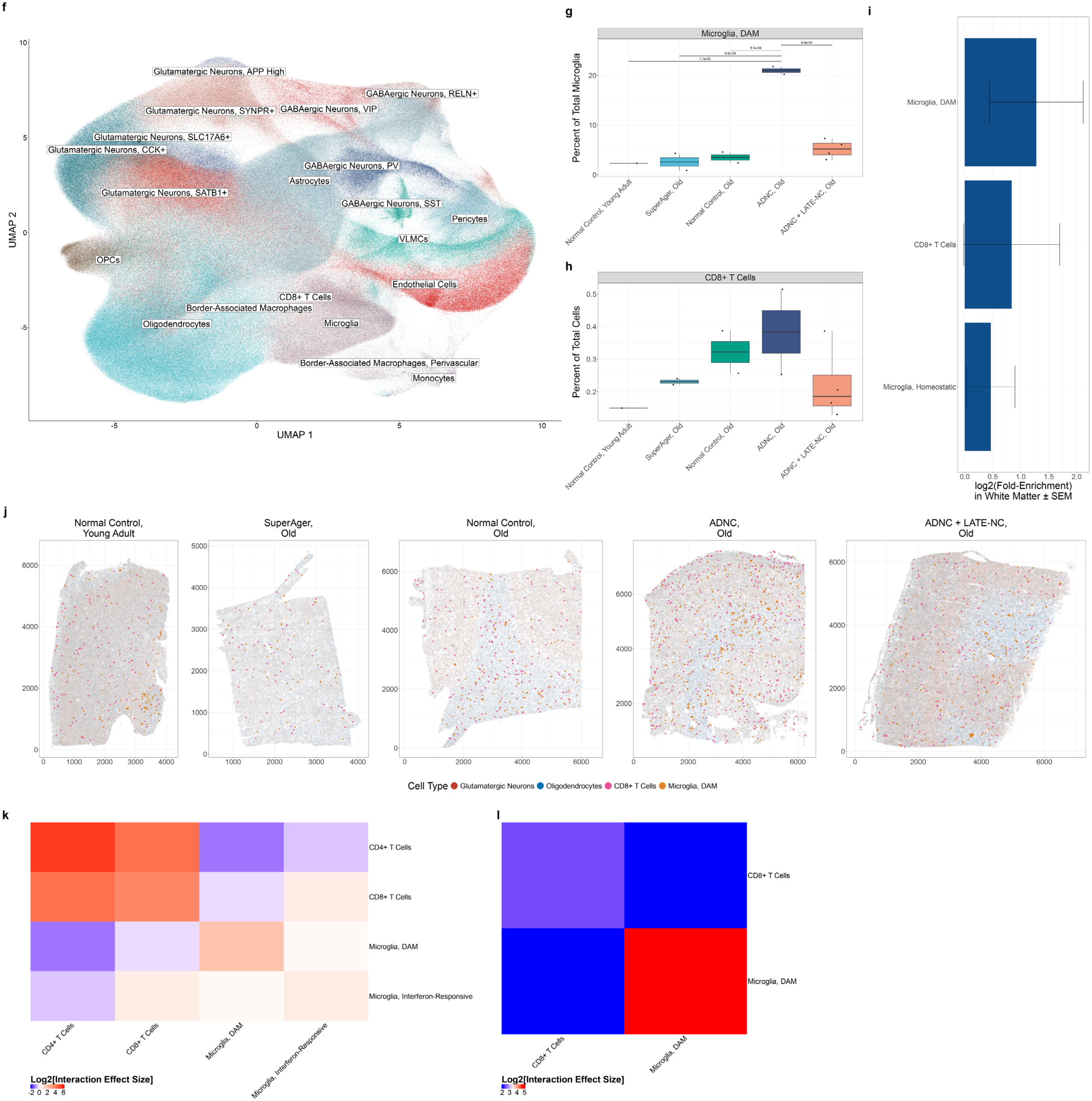
Spatial distributions of age-associated neuroimmune cell states in mice and humans. **(a)** UMAP of imaging spatial transcriptomics results from 2,971,436 cells passing quality filters segmented from whole mouse skulls of young adult (4mo; n = 2) and old (18mo; n = 2) C57BL/6JN mice. **(b)** Boxplots of DAM and IRM abundance in young vs. old animals from (a) as a percent of total microglia. Comparisons are t-tests with FDR correction. **(c)** Boxplots of CD8+ T cell abundance in young vs. old animals from (a) as a percent of total cells. Comparisons are t-tests with FDR correction. **(d)** Log2 fold-enrichment of select microglia and T cell states in white matter as compared to a random distribution for each mouse in (a). Only parenchymal cells were included in analysis. **(e)** Spatial distributions of DAM (orange), IRM (green), and CD8+ T cells (pink) from (a). White matter regions are delineated by oligodendrocytes (transparent blue), and gray matter regions are delineated by glutamatergic neurons (transparent red). **(f)** UMAP of imaging spatial transcriptomics results from 1,360,254 cells passing quality filters segmented from middle frontal gyrus (MFG) brain sections from 11 human patients postmortem, consisting of a young adult control (28yo; n_male_ = 1); normal, elderly controls (87-92yo; n_male_ = 1, n_female_ = 1); SuperAgers (85-91yo; n_male_ = 1, n_female_ = 1); amnestic dementia (AD) with ADNC pathology (62-66yo; n_male_ = 2); and amnestic dementia (AD) with ADNC + LATE-NC pathology (75-97yo; n_female_ = 3; n_male_ = 1). **(g)** Boxplots of DAM abundance in human postmortem MFG samples from (f) as a percent of total microglia. Comparisons are t-tests with FDR correction. **(h)** Boxplots of CD8+ T cell abundance from (f) as a percent of total cells. **(i)** Log2 fold-enrichment of select microglia and lymphocyte cell states in white matter as compared to a random distribution for each patient in (e). **(j)** Spatial distributions of DAM (orange) and CD8+ T cells (pink) from (f). White matter regions are delineated by oligodendrocytes (transparent blue), and gray matter regions are delineated by glutamatergic neurons (transparent red). **(k-l)** Spatial correlation between select cell states in whole mouse skulls (k) and postmortem human MFG (l). All correlations shown are significant (q < 0.05, global envelope test against a null hypothesis of random cell state distributions with FDR correction).

To credential these findings in human aging, we profiled sections of human middle frontal gyrus from postmortem brains by imaging spatial transcriptomics. We included brain samples from young adult research participants, older participants with typical or “average” cognition based on normative samples, and participants diagnosed with amnestic dementia representing a pathological aging trajectory. To explore the full range of successful aging, which encapsulates average and better-than-average cognitive phenotypes, we also included unique participants defined as SuperAgers, individuals with memory function equivalent to individuals 20-30 years their junior (Supp. Table 4)^50^. We identified populations of excitatory and inhibitory neurons, oligodendrocytes, oligodendrocyte precursor cells, astrocytes, microglia, BAMs endothelial cells, vascular leptomeningeal cells, CD8+ T cells, and monocytes (Fig 2f; Extended Data 3). We identified human DAM, characterized by expression of *SPP1*, *LPL*, *GPNMB*, *CST7*, and *CSF1*; and CD8+ T cells expressing *CD8A*, and *CD3E* (Supp. Fig. 2f-h). Recognizing the limitations of sampling, we observed qualitative increases in both DAM and CD8+ T cells during normal and pathological aging, particularly in patients diagnosed with amnestic dementia with Alzheimer’s disease neuropathic change pathology (ADNC)^51–53^. Curiously, these associations were diminished in individuals with comorbid ADNC and limbic-predominant age-related TDP-43 encephalopathy neuropathic change (LATE-NC)^54^. SuperAgers exhibited a qualitative reduction in both DAM and CD8+ T cell abundance, suggesting that DAM are associated specifically with more pathological aging trajectories (Fig. 2g-h)^55^. Mirroring our findings in mice, DAM and CD8+ T cells were modestly enriched in white matter tracts underlying the middle frontal gyrus (MFG) in our samples, likely sampled from short association fibers (U-fibers) that comprise portions of the superior longitudinal fasciculus and inferior fronto-occipital fasciculus (Fig. 2i-j; Supp. Fig. 2i). In both human and mouse brains, DAM clustered together and exhibited strong significant spatial autocorrelation, suggesting a response to a local stimulus in the CNS microenvironment (Fig. 2k,l; Supp. Fig. 2j,k). Collectively, these findings further establish white matter regions as centers of age-related neuroinflammation in both mice and humans.

### Influenza A virus pneumonia accelerates age-related DAM accumulation and activates T cells

We used our single cell object to quantify cell-state changes in microglia during the acute phase of IAV pneumonia and after recovery (Fig. 3a). Notably, not a single count of IAV RNA was observed in any cell in the dataset, even prior to gene filtering, strongly arguing against CNS infection (Supp. Fig. 3a). In young adult mice, the abundance of IRM, DAM, CD4+ T cells, and CD8+ T cells did not change over the course of IAV pneumonia and recovery. In middle-aged mice, however, DAM increased in abundance during acute IAV pneumonia and persisted after recovery at levels that were similar to those observed in old, naïve mice. In old mice, the abundance of IRMs and DAM was increased prior to IAV pneumonia, relative to young adult mice, and did not change significantly over the course of IAV pneumonia and recovery. Despite ongoing invasion of T cells into the brain during normal aging (Fig 1h,j), these numbers were not significantly altered over the course of IAV pneumonia and recovery (Supp. Fig. 3b-c). Rather, existing resident CD4+ and CD8+ T cells exhibited concerted shifts toward aging-associated populations in middle-aged mice. CD4+ T cells transiently shifted toward a TCM state, while CD8+ T cells shifted toward a TEM state that persisted after recovery (Fig. 3c). Transient increases in IRM and interferon-responsive CD8+ T cells were also observed during acute IAV pneumonia in middle-aged animals, however these cell states returned to middle-age, naïve levels after recovery (Fig 3b; Supp. Fig. 3d). No novel cell states emerged during either acute IAV pneumonia or after recovery in any age group.

**Figure 3.**
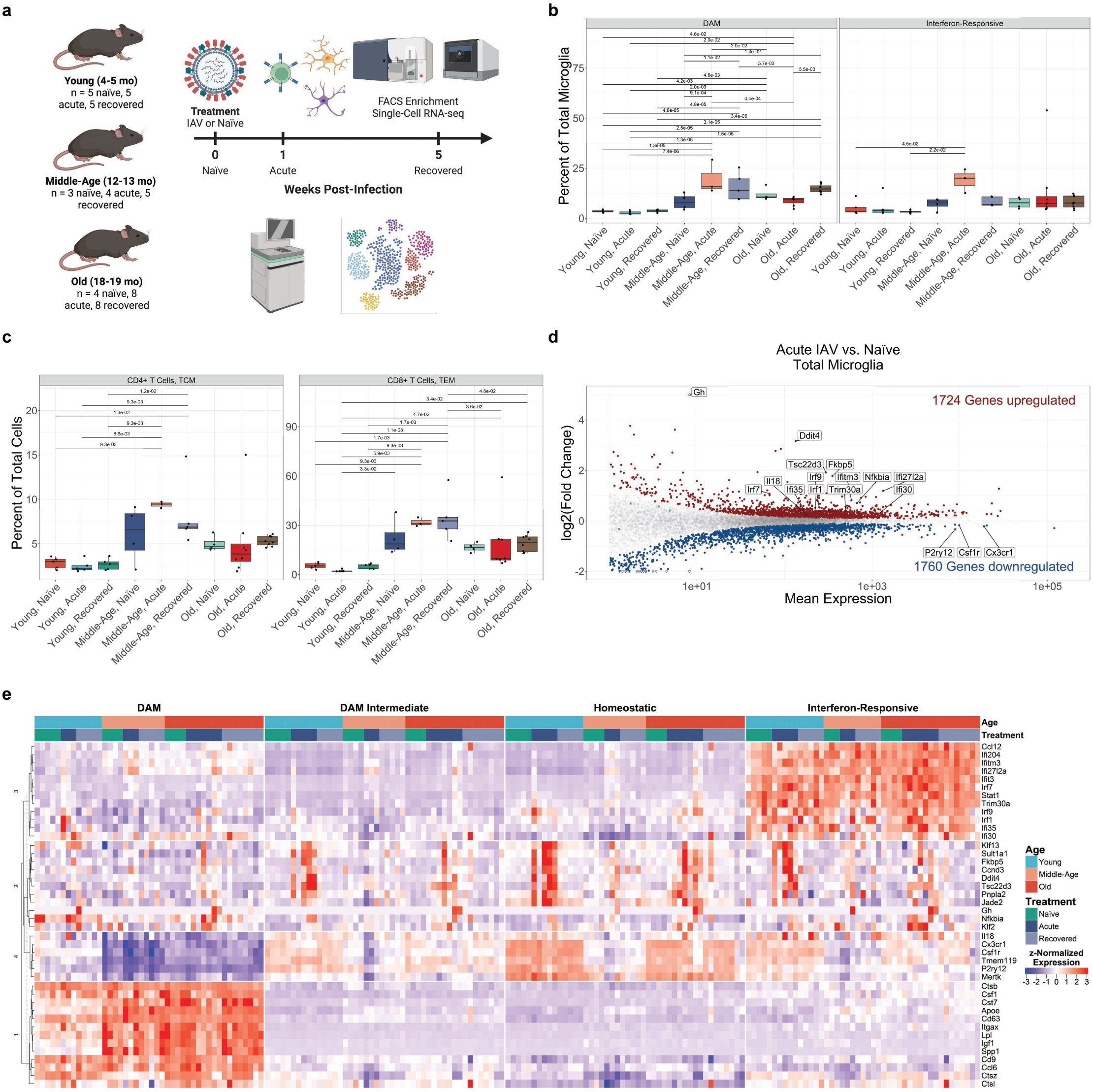
Influenza A pneumonia accelerates the formation of DAM in middle aged mice. **(a)** Experimental design and timeline of aging and viral pneumonia experiments. **(b)** Abundances of DAM and IRM from scRNA-seq data as a proportion of total microglia from whole brains of C57BL/6JN mice in the steady state (naïve), during acute IAV infection (1 weeks post-infection), and after recovery from pneumonia (5 weeks post-infection) in young adult (4-5mo; n_naïve_ = 5, n_acute_ = 5, n_recovered_ = 5), middle-aged (12-13mo; n_naïve_ = 3, n_acute_ = 4, n_recovered_ = 5), and old age (18-19mo; n_naïve_ = 4, n_acute_ = 8, n_recovered_ = 8). Comparisons are t-tests with FDR correction. **(c)** Abundances of CD4+ and CD8+ T cell states from scRNA-seq data a proportion of total cells. **(d)** MA plot of pseudo-bulk RNA-seq data comparing microglial transcriptomic responses to IAV pneumonia, irrespective of age. Significantly (q < 0.05, Wald test) upregulated genes are shown in red, and significantly downregulated genes are shown in blue. Genes shown in gray are not significantly differentially expressed. Genes represented by triangles are outside the plot limits. **(e)** Hierarchical clustering of mean gene expression in microglia by cluster for select genes responsive to acute IAV (Module 2) and markers of DAM (Module 1), IRM (Module 3), and homeostatic microglia (Module 4). Modules were split using k-means clustering with k = 4.

To further characterize these results at the molecular level, we identified differentially expressed genes in middle-aged animals compared to young adult animals during acute IAV infection by pseudo-bulk differential expression analysis of total microglia. Microglia from all age groups exhibited a core set of IAV response genes during acute IAV pneumonia, characterized by anti-inflammatory genes involved in glucocorticoid signaling, including *Gh*, *Tsc22d3*^56^, *Fkbp5*^57^, *Pnpla2*^58^, and *Ddit4*^59^, interferon-responsive gene expression, and expression of the cell-cycle regulator *Ccnd3*, which returned to baseline after recovery (Fig. 3d,e)^60,61^. Sub-cluster analysis of microglia revealed that all microglia clusters participated in the “core” IAV response (Module 2), however changes in DAM marker gene expression (Module 1) were driven by changes in microglia composition, rather than gene expression changes in existing clusters, with the exception of upregulation of *Apoe* in middle-aged animals in response to IAV (Fig. 3e). Pseudobulk analysis of only homeostatic microglia from all three age groups confirmed these findings (Supp. Fig. 3e-g). The most significant effect of IAV pneumonia was therefore an acceleration of DAM formation in middle-aged animals to levels of their old counterparts, some 6 months prematurely. As DAM did not increase in homeostatic microglia during IAV irrespective of age, these findings suggest the expansion of DAM we observe in response to influenza results from local injury to neurons or other cells in the CNS rather than a cell-autonomous response to influenza across microglia.

### Age-related changes in microglia cell states are driven by the aging brain microenvironment

We wondered whether the age-related cell state shifts in microglia are cell-intrinsic, or if they are driven by the aging brain microenvironment. Microglia rely on tonic activation of CSF1R for their proliferation and survival. The administration of pharmacological inhibitors of CSF1R depletes microglia, which are repopulated after drug discontinuation from spatially restricted regions of the CNS^62^. We reasoned that if the age-related transcriptomic changes in microglia resulted from the direct effect of inflammatory cytokines or other signals originating outside the CNS they would be reversed in repopulated microglia after their depletion. In contrast, if these endocrine signals targeted other CNS populations that resulted in secondary changes in microglia they would not. We therefore ablated microglia with the CSF1R antagonist PLX3397 and profiled flow-cytometry sorted microglia by bulk RNA-seq in the steady state (control chow), after 14 days of PLX3397 treatment, and 7 and 28 days after PLX3397 discontinuation.^63^ We did not observe reductions in DAM or IRM gene marker expression phenotype in old animals after 7 or 28 days of repopulation and reintegration into the CNS microenvironment (Fig. 4b-c; Supp. Fig. 4b; Extended Data 4), suggesting that the aging brain microenvironment drives cell-state shifts in microglia.

**Figure 4.**
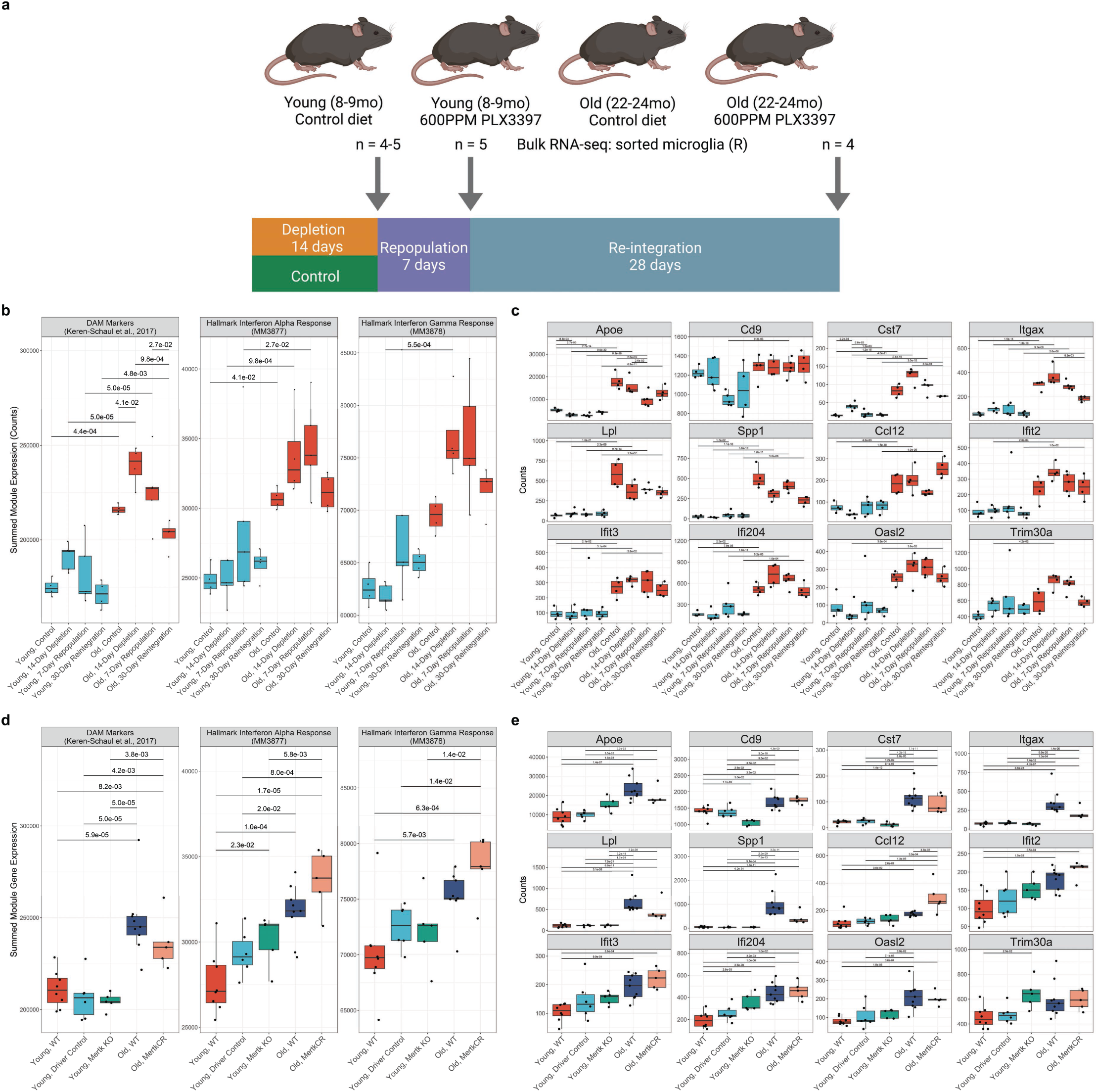
Aging drives the development of DAM and IRM through non-cell-intrinsic mechanisms. **(a)** Visual methods for microglial depletion and repopulation experiments for control chow: young adult (8mo; n = 4), old (23mo; n = 4); 14-day depletion with 600ppm PLX3397 chow: young adult (n = 4; 8mo), old (n = 4; 22mo), 7 days after PLX3397 withdrawal: young adult (n = 5; 8mo), old (n = 5; 21mo), and 30 days after PLX3397 withdrawal: young adult (n = 4; 8mo), old (n = 4; 22mo). **(b)** Summed modular expression of DAM marker genes^22^ and interferon-responsive marker genes from RNA-seq performed on flow-cytometry sorted microglia. Comparisons are Wilcoxon rank sum tests with FDR correction. **(c)** Expression of select DAM and IRM marker genes from RNA-seq performed on flow-cytometry sorted microglia from (b). Comparisons are Wald tests with FDR correction in DESeq2. **(d)** Summed modular expression of DAM marker genes and interferon-responsive marker genes from RNA-seq performed on flow-cytometry sorted microglia from young adult, wild-type mice (n = 8; 5-7mo; reused from 1A); young adult driver controls (n = 6; 3-6mo); young adult mice with 30 or 60-day conditional deletion of *Mertk* in microglia (n = 5; 7-8mo); old, wild-type mice (n = 9; 20-21mo; reused from 1A); and old mice with a cleavage-resistant *Mertk* variant (n = 5; 17mo).^66^ No significant differences were observed between 30 and 60 day time-points and these results were subsequently binned together. Comparisons are Wilcoxon rank sum tests with FDR correction. **(e)** Expression of select DAM and IRM marker genes from RNA-seq performed on flow-cytometry sorted microglia from (d). Comparisons are Wald tests with FDR correction in DESeq2.

Phagocytosis of damaged cells in the CNS is a major function of microglia. Indeed, *Gpnmb*, which was recently shown to contribute to the phagocytic function of microglia, was among the most upregulated genes with aging in our bulk RNA sequencing dataset and its expression is localized to DAM. The tyrosine kinase receptor MERTK interacts with its ligand Gas6 and Protein S to phagocytose phosphatidylserine residues on the outer surface of damaged or apoptotic cells in the CNS^63,64^. Fatty acid beta oxidation following efferocytosis has been shown to be net anti-inflammatory in macrophages^65,66^ and loss of *Mertk* has been shown to reduce lifespan in AD models.^67^ Exposure to apoptotic bodies derived from neurons has also been shown to be sufficient to induce a DAM-like state in human iPSC-derived microglia.^68^ Furthermore, we observed significant downregulation of *Mertk* in microglia during physiological aging (Fig. 1a). We therefore wondered whether reduced *Mertk* expression in microglia is sufficient to drive premature DAM and/or IRM expansion. We performed bulk RNA-seq on flow-cytometry sorted microglia from young adult *Cx3cr1-CreERT2; zsGreen^lsl/-^ x Mertk^flox/flox^* mice and driver controls 30 or 60 days after tamoxifen administration and compared DAM and IRM marker gene expression (Supp. Fig. 4c; Supp. Table S5). We did not observe expansion of DAM or IRM in young adult mice in the absence of *Mertk*, supporting previous findings in the developing retina^69^. MERTK is selectively cleaved from the surface of macrophages during inflammation, leading to reductions in phagocytosis^66^. We reported that mice expressing a cleavage-resistant variant of MERTK are resistant to some age-related phenotypes^48,70^. We therefore reasoned that expression of cleavage-resistant MERTK [MERTK(CR)] might reduce age-related neuroinflammation. We did not observe changes in the expression of DAM and IRM marker genes in microglia purified by flow cytometry from old mice expressing MERTK-CR throughout their lifespan and old, wild-type mice (Fig. 4d,e; Extended Data 5). These results suggest that prolonged expression of functional MERTK in aging is not sufficient to rescue age-related expansion of DAM and IRM. Collectively, these findings suggest that age-related shifts in microglial cell state are driven primarily by the aging CNS microenvironment and are independent of MERTK.

## Discussion

We used unbiased bulk and single-cell transcriptomics, flow cytometry, metabolomics, and imaging spatial transcriptomics to profile neuroimmune cells in the course of normal aging and in response to IAV pneumonia. We show that IRM and DAM expand beginning in middle age, and peak in old age. With a similar cadence, several CD8+ and CD4+ T cell populations, including TEM, infiltrate the brain. The age-related expansion of DAM is accompanied by broad remodeling of microglial metabolism, including aerobic glycolysis, citrulline depletion, and an increased ratio of itaconate/α-ketoglutarate, correlated with a broken TCA cycle. In mice, DAM and CD8+ T cells were localized to white matter regions. In human brains, DAM were similarly localized to white matter, where their abundance was qualitatively associated with aging and strongly associated with ADNC pathology. Curiously, however, this association was not observed in the presence of comorbid ADNC and LATE-NC pathology, a valuable topic for further investigation. In both human and mouse brain tissue, DAM clustered tightly together, suggesting a response to focal pathology. Immunosenescent-like signatures were localized to cell states expanded during aging, including DAM and CD8+ T cells.

Irrespective of age, microglia responded to IAV pneumonia by activating a core set of glucocorticoid response genes and the cell-cycle regulator *Ccnd3*, likely reflective of pro–resolution, anti-inflammatory signaling. These changes developed in the absence of direct CNS infection. In middle-aged animals, these changes were accompanied by an expansion of DAM and CD8+ TEM to levels similar to old animals that persisted after recovery from IAV pneumonia. This 6-month difference in mice is the equivalent of ∼30 years in humans.^34^ In old animals, the abundance of DAM was increased even before IAV pneumonia and did not increase significantly in response. Whether this represents a ceiling on the formation of DAM in the absence of overt pathology or a change in CNS susceptibility to viral pneumonia with advancing age is not known.

Microglia are long lived cells that originate from the fetal yolk sac^71,72^. Some have therefore suggested that microglia undergo cell-autonomous changes during aging which prime them to differentially respond to systemic inflammation^28–30,73^. While neuroinflammation was observed in old (and to a lesser degree middle-aged) animals, the neuroinflammatory response induced by acute IAV was similar in all age groups, arguing against age-associated pro-inflammatory “priming” in old animals. Alternatively, Jurgens and colleagues showed that influenza infection resulted in changes to neuronal structure, which might suggest changes in microglia after influenza are responsive to changes in the CNS microenvironment^11^. Accordingly, we transiently depleted microglia from young adult and old mice using the CSF1R antagonist PLX3397 and allowed them to repopulate. The lack of change in age-related microglia phenotypes upon repopulation suggests that the microenvironment is substantially responsible for age-related changes in microglia. Our experiments with mice harboring gain and loss of function mutations in the tyrosine kinase receptor MERTK are consistent with these findings. MERTK is important for the phagocytosis of cells that have undergone apoptosis (efferocytosis) and for the pruning of synapses during neuronal development (trogocytosis)^74,75^. We found that, although *Mertk* expression was reduced in microglia from old mice, loss of MERTK in young adult mice did not cause a premature expansion of DAM and/or IRM. Conversely, old mice harboring a gain of function in MERTK were not protected against the age-related expansion of DAM. Our findings are consistent with other groups, who reported that age-related DAM expansion is unaffected by heterochronic parabiosis and that myeloid cells adoptively transferred into the brains of old animals assume a DAM phenotype.^76,77^

In conclusion, we provide a comprehensive molecular and cellular neuroimmune atlas of the aging mouse brain during homeostasis, non-neuroinvasive pneumonia, and recovery. We identify white matter expansion of DAM and IRM and invasion of effector CD8+ T cells as hallmarks of physiological brain aging. Our studies suggest a point of CNS vulnerability to IAV pneumonia in middle age, when the ability to resolve changes in the microenvironment induced by pneumonia is lost. Guided by the spatial localization of DAM to white matter regions, future studies should focus on understanding regional neuropathology induced by IAV pneumonia, perhaps changes to myelin, that drive DAM phenotypes.

There were limitations to our study. First, we limited our study to male mice. While this is partially justified by the worsened outcomes after viral pneumonia in men compared with women, a finding recapitulated in mice, this limitation was largely pragmatic and precludes our exploration of sex-specific differences in the microglial response to influenza. Second, while our findings implicate changes in the microenvironment as drivers of the DAM phenotype, depletion of microglia with PLX3397 is incomplete and repopulation occurs from CNS progenitors. Hence, we cannot exclude cell-autonomous changes in microglia remaining after PLX3397 depletion or in microglia progenitors as drivers of the age-related expansion of DAM. Third, we were not able to associate changes in the microglial response to IAV pneumonia with aging with cognitive performance. Common cognitive tests in mice reliably detect the age-related decline in cognitive function associated with normal aging and IAV pneumonia in relatively small cohorts of mice. However, old animals have a lower baseline and more variable performance on these tests. Hence, detecting the decline and recovery of cognitive function over the course of IAV pneumonia with sufficient statistical power in old mice would require hundreds of animals. More sensitive markers of behavior, reflective of CNS pathology adapted to aging will be needed to address these questions. Finally, while our studies exclude MERTK as sufficient or necessary for the age-related expansion of DAM with advancing age, other molecules important for microglia function might synergize with changes in the microenvironment to drive age-related DAM expansion.

## Extended Data

**Extended Data 1**. Differential expression analysis (DEA) results from bulk RNA-seq comparing young vs. old mice (related to Fig. 1a).

**Extended Data 2.** Marker genes for mouse scRNA-seq clusters (related to Figs. 1 and 3) and pairwise differential expression analysis (DEA) results from pseudobulk RNA-seq for all cell types and groups in mouse scRNA-seq data (related to Fig. 3).

**Extended Data 3.** Marker genes for mouse and human imaging spatial transcriptomics clusters (related to Fig. 2) and mouse and human 10X genomics Xenium panel genes.

**Extended Data 4**. Pairwise differential expression analysis (DEA) results, and k-means gene assignment from bulk RNA-seq of microglia from depletion and repopulation experiments during aging.

**Extended Data 5.** Pairwise differential expression analysis (DEA) results from bulk RNA-seq of microglia with loss- or gain-of-function mutations in MERTK in aging.

## Materials and Methods

### Sex as a biological variable

For human studies, samples from both male and female patients were included. The costs of scRNA-seq and spatial transcriptomics only allowed us to study one sex. Because viral pneumonia disproportionately impacts men, we chose to include only male mice in all mouse experiments.

### Mice

All experimental protocols were approved by the Institutional Animal Care and Use Committee at Northwestern University. All strains including wild-type mice are bred and housed at a barrier- and specific pathogen-free facility at the Center for Comparative Medicine at Northwestern University. We used definitions for murine age from the Jackson Laboratory, which defines mature adult / young adult mice as 3-6 months old, middle aged mice as 10-14 months old, and old mice as 18-24 months of age and very old mice as >24 months of age^34^. Young adult mice were used from 4-6mo, and old mice were 18-21mo for all experiments. All wild-type mice were obtained from the National Institute of Aging. The Cx3cr1-CreERT2 mice,^78^ zs-Green^lsl^,^79^ mice were obtained from Jackson Laboratories (Jax stocks 020940, and 007906, respectively). Mertk^flox/flox^ mice were a gift from Shelton Earp.^80^ Mertk-CR mice were a gift from Ira Tabas via Edward Thorp.^66^

### Influenza A virus culturing and propagation

Influenza A virus (A/WSN/1933 [H1N1]) was cultured and propagated as described previously.^81^ Briefly, Influenza virus strain A/WSN/1933 (WSN) was grown for 48 h at 37.5°C and 50% humidity in the allantoic cavities of 10–11-day-old fertile chicken eggs (Sunnyside Hatchery). Viral titers were measured by plaque assay in Madin-Darby canine kidney epithelial cells (American Type Culture Collection). Stocks were frozen in liquid nitrogen and stored for up to one year. Working aliquots were stored at −80°C and underwent a total of two freeze-thaw cycles on ice.

### Intratracheal instillation of influenza A virus

Mice were infected with mouse-adapted influenza A virus (A/WSN/1933 [H1N1]) as described previously^82^. Briefly, mice were anesthetized with isoflurane and intubated using a 20 gauge angiocath. Young adult mice were then instilled with 20 plaque-forming units (pfu) and old mice were instilled with 5 pfu of mouse-adapted influenza A/WSN/33 [H1N1] virus in 50 µL of ice-cold PBS through the catheter. Naïve/control mice were left untreated. Mice were then monitored every 1-2 days for weight loss and mortality until their respective endpoints.

### Oral gavage of tamoxifen

For induction of Cre-ERT2 activity, 10 mg tamoxifen (Sigma, T5648) was dissolved in 100µL corn oil (Sigma, C8267) and administered to the anesthetized mice by oral gavage. Mice were treated once per day for 2 days. Mice were monitored for weight loss for at least 1 week after the second dose. No mice showed overt weight loss. *Cx3cr1-CreERT2* x *zsGreen^lsl^* mice were euthanized 16 days-post-treatment after monocyte turnover. *Cx3cr1-CreERT2; zsGreen^lsl^* x *Mertk^flox/flox^* mice were euthanized 16 or 61 days-post-treatment.

### Microglia depletion with PLX3397

Mice were fed either Teklad LM-485 control chow alone (Envigo 7012) or control chow laden with 600ppm PLX3397 (Medkoo 206178) for 14 days *ad libitum*. Microglia were then isolated for bulk RNA-seq at 0, 7, or 28 days after return to control chow as described in^31^. Bedding was changed at each chow change to prevent delayed or prolonged drug treatment via coprophagy.

### Brain tissue processing and flow cytometry isolation of neuro-immune cells

Mice were euthanized by IP injection with 250µL of a 1:10 solution of Euthasol (Virbac) in PBS. After loss of paw withdrawal reflex, mice were perfused transcardially with 10mL ice-hold HBSS followed by decapitation. Brains were removed and split by hemisphere. For bulk RNA-seq, right hemispheres were fixed for 24-48hrs in ice-cold 4% PFA before transfer to PBS + 0.01% sodium azide indefinitely at 4°C. Left hemispheres were transferred 0.5mL ice-cold digest buffer consisting of 1X Papain Dissociation System (Fisher NC9212788; 1 vial dissolved in 5mL HBSS to yield 20U of papain/mL in 1mM L-cysteine with 0.5mM EDTA) and 1mg/mL DNAse I (Roche 10104159001) with curved scissors. For scRNA-seq, both hemispheres were used for dissociation. Chopped tissue was then transferred to gentleMACS C-tubes (Miltenyi 130-093-237) and mixed with 1mL HBSS. Samples were then mechanically dissociated using a gentleMACS Octo Dissociator using the stock program “m_brain_03_01”. Samples were then shaken at 200rpm, 37°C for 30 minutes, followed by a second round of mechanical dissociation. Digestion was then stopped by mixing samples with 9mL ice-cold, sterile-filtered 1% BSA (Sigma SLBW2268) in 1X HBSS. Cell suspensions were then mashed through a 70µm filter with 4×10mL ice-cold 1% BSA in HBSS into a fresh 50mL conical tube. The resulting single-cell suspension was then pelleted at 400 x *g* for 10 minutes at 4°C and resuspended in 25mL ice-cold 40% Percoll (Millipore-Sigma GE17-0891-01) in 1X HBSS without calcium or magnesium (Gibco 14185-052) and pelleted at 500 x *g* for 30 minutes at 4°C. The myelin-debris-containing supernatant was then removed using a vacuum pipet and the remaining <1mL was adjusted to 10mL with ice-cold HBSS, spun down at 400 x *g* for 5 minutes at 4°C, and resuspended in 50µL 1:50 mouse Fc block (BD Pharmingen 553142; clone 2.4G2) in MACS buffer and incubated for a minimum of 5 minutes on ice. A 2µL aliquot was then counted on a Cellometer K2 as above. For microglia bulk RNA-seq, 50µL of the antibody cocktail shown in Supplementary Table S4 was added to the existing suspension. For scRNA-seq of mixed neuro-immune cells, 50µL of the antibody cocktail shown in Supplementary Table S5 was added, including hashtagging antibodies. Suspensions were mixed by pipetting and incubated for 30 minutes at 4°C in the dark. Suspensions were then mixed with 900µL of MACS buffer, pelleted at 400 x *g* for 5 minutes at 4°C, and resuspended in 300-800µL of MACS buffer based on cell density. Immediately prior to sorting, cells were again filtered Immediately before sorting, suspensions were filtered through a 70µm filter, rinsed with 200µL MACS buffer. SYTOX stain was then added at 1µL/mL suspension and the suspension was mixed thoroughly. For bulk RNA-seq, microglia were sorted on a FACS Aria II (BD Biosciences) as non-debris, live (SYTOX^−^), CD45^int^, Ly6C^−^, [CD3e/NK1.1/Ly6G]^−^, CD64^+^, CD11b^+^, F4/80^+^, MHC-II^+/−^ events directly into 100µL PicoPure lysis buffer (Arcturus KIT0204) at 7°C with occasional mixing to prevent layering (Fig. S5). For Cx3cr1-CreERT2 x zsGreen^lsl^ mice, zsGreen was used in place of F4/80 (Fig. S6). After sorting a maximum of 80k cells, the solution was homogenized by vigorous pipetting and placed on ice until the completion of sorting. All tubes were then transferred to a −80°C freezer until RNA isolation. For scRNA-seq of neuro-immune cells, all cells were sorted using a FACS ARIA II (BD Biosciences) or FACSymphony S6 (BD Biosciences) into a single tube with 400µL of MACS buffer. Microglia were sorted as non-debris, live (SYTOX^−^), CD45^int^, CD3e^−^, CD11b^+^, CD64^+^, [NK1.1/Ly6G/Ly6C]^−^ events. T cells were sorted as non-debris, live (SYTOX^−^), CD45^hi^, CD3e^+^ events. Astrocytes were sorted as non-debris, live (SYTOX^−^), CD45^−^, CD3e^−^, ACSA-2^+^ events. A total of 100k-120k cells were sorted for all samples. For bulk metabolomics on sorted microglia, microglia were sorted on a FACSymphony S6 (BD Biosciences) as non-debris, live (SYTOX-), CD45int, Ly6C-, [NK1.1/Ly6G]-, CD3e-, CD64+, CD11b+, CD206-events into MACS buffer. Cells were then pelleted and resuspended in 80% LCMS-grader acetonitrile (Supelco 1.00029.1000) in LCMS-grade water (Supelco 1.15333.1000) at 2-5×10^3^ cells/µL followed by lysis with trituration and vigorous vortexing. Lysates were then snap-frozen in liquid nitrogen. Prior to metabolomics analysis, cells were thawed on ice and centrifuged at 17,000 x g for 30 minutes at 4°C. Supernatants were then transferred to fresh tubes for analysis.

### Bulk RNA-seq

Frozen cell lysates were first incubated at 42°C for 30 minutes in a water bath to thaw and ensure complete lysis. RNA was then isolated using the Arcturus PicoPure RNA extraction kit (Arcturus KIT0204) according to manufacturer’s instructions. RNA quality and quantity were assessed using TapeStation 4200 High Sensitivity RNA tapes (Agilent), and RNA-seq libraries were prepared from 850-2056pg of total RNA using SMARTer Stranded Total RNA-seq Kit v2 (Takara Bio). After QC using TapeStation 4200 High Sensitivity DNA tapes (Agilent), dual-indexed libraries were pooled and sequenced on a NextSeq 500 instrument (Illumina) for 75 cycles, single-end. FASTQ files were generated using bcl2fastq 2.20 (Illumina) using default parameters. To facilitate reproducible analysis, samples were processed using the publicly available nf-core/RNA-seq pipeline version 3.5 implemented in Nextflow 22.04.5.5708 using Singularity 3.8.1 with the minimal command nextflow run nf-core/rnaseq \ -r ‘3.5’ \ -profile nu_genomics \--additional_fasta ‘transgenes.fa’ \ --star_index false \ --three_prime_clip_r2 3 \ --genome ‘GRCm38’. All analysis was performed using custom scripts in R version 4.1.1 using the DESeq2 version 1.34.0 framework^83^. A “local” model of gene dispersion was employed as this better fit dispersion trends without obvious overfitting, and pairwise comparisons were performed on a combined factor of age, treatment, and genotype. Alpha was set at 0.05 for all DEA. Otherwise default settings were used. See code for details. High-level analysis was performed using custom scripts available in the nu-pulmonary/utils GitHub repository.

### Single-cell RNA-seq

Sorted cells were diluted to 1.5mL with BamBanker medium and immediately pelleted at 400 x *g* for 5 minutes at 4°C. Cells were then resuspended in BamBanker at ∼1×10^6^ cells/mL. Cell concentrations and viability were confirmed using a Cellometer K2 as above. Libraries were then generated using the 10X Genomics 3’ V3.1 kit, according to manufacturer’s instructions (CG000317 Rev C), using a 10X Genomics Chromium X Controller. After quality checks, single-cell RNA-seq libraries were pooled and sequenced on an Illumina NovaSeq 6000 instrument. CITE-seq (hashtagging) libraries were sequenced independently on an Illumina NextSeq 2000 instrument.

### Single-cell RNA-seq Analysis and Processing

Data were processed using the Cell Ranger 7.0.1 pipeline (10x Genomics) with intronic reads disabled. To enable detection of viral RNA, reads were aligned to a custom hybrid genome containing GRCm38.93 and Influenza A (GCF_001343785.1). Putative heterotypic doublets were then flagged for removal using scrublet 0.2.3 with manual thresholding of doublet scores, before removal using custom scripts in R^84^. Cell calling was performed using the cellranger pipeline. Thresholding of Initial filtering and preprocessing was performed using Seurat 4.4.0^85^, followed by integration using SCVI within SCVItools 0.14.3^86^, and re-imported into Seurat for all clustering, dimensional reduction, and all downstream high-level analysis using inbuilt Seurat functions and custom scripts in R. All manipulations in Seurat were performed with the aid of tidyseurat 0.8.0^87^. Normalization was performed using SCTransform 0.4.1^88^, and clustering was performed using the Leiden algorithm. Default parameters were otherwise used unless directly specified^89^. Module scores for select gene sets were calculated using the AddModuleScore() function in Seurat using default parameters. Median scores were then calculated and reported for each cell state (e.g. DAM, CD8+ TEM) for each biological replicate (e.g. mouse, patient).

### Pseudobulk RNA-seq

Pseudobulk analysis was performed using the “bulkDEA” function in the “Seurat_pseudobulk_DEA.R” script in the NUPulmonary/utils repository. Briefly, raw counts (object@assays$RNA@counts) were aggregated by sum by sample (e.g. mouse or patient) and major cell type (e.g. microglia, CD8+ T cells) or cell state (e.g. DAM) as indicated and passed from Seurat to DESeq2 1.34.0 ^83^ with relevant metadata. Samples with fewer than 20 cells for a given comparison cell type or state were excluded from analysis. Size factor estimation, dispersion fitting, and Wald tests were performed using the DESeq function in DESeq2. “Parametric” and “local” models of dispersion were compared visually for goodness-of-fit, and the most reasonable fit was chosen. Results were then extracted using the results function with alpha set to 0.05. For mouse data, DEA was performed across a combined factor of age and IAV treatment group, e.g. old_acute vs young_acute. In order to determine the “core” response to IAV pneumonia, irrespective of age, gene expression changes were modeled with the design formula ∼ flu_group + age_group, and only the flu_group contrast was considered (acute vs naïve). Default parameters were used unless otherwise specified. In all plots of pseudobulk gene counts, p-values shown are FDR-corrected p-values directly from DESeq2 analysis.

### Clinical Characterization

Four participants had an antemortem clinical diagnosis of amnestic dementia based on National Institute on Aging–Alzheimer’s Association (NIA-AA) criteria^53^. Four participants met criteria for cognitive SuperAger status based on outstanding memory performance prior to death. Episodic memory was assessed using the delayed recall score of the Rey Auditory Verbal Learning Test (RAVLT)^90^. SuperAgers were required to perform at or above average for individuals in their 50s–60s, defined as a RAVLT delayed recall raw score ≥9 and scaled score ≥10 (see Gefen et al., 2015, for further details)^91^. For comparison, two cognitively average, older adults (“Normal Controls”) and one younger adult (“Young Control”) were identified based on extensive chart review and neuropsychological assessment if available. Data on whether participants suffered from infection requiring hospitalization (<1 year prior to death) were noted. See Supp. Table 4.

### Neuropathological Evaluation and Histological Preparation of Human Tissue

Autopsied brains from 11 participants were acquired from the Northwestern University Alzheimer’s Disease Research Center (ADRC) Brain Bank. Written informed consents were obtained from all participants, and the study was approved by the Northwestern University Institutional Review Board and in accordance with the Helsinki Declaration. Following autopsy, the cerebral hemispheres were separated in the midsagittal plane, cut into 3- to 4-cm coronal slabs, fixed in 10% formalin for 2 weeks or 4% paraformaldehyde for 36 hours, taken through sucrose gradients (10%–40%) for cryoprotection, and stored in 40% sucrose at 4° C. Samples of the middle frontal gyrus (MFG) were embedded in paraffin. Neuropathological diagnoses were rendered based on published consensus criteria^51,52,54^.

### Imaging spatial transcriptomics

For mouse samples, young (4mo) and old (18mo) male mice were euthanized with 150µL of 1:10 diluted Euthasol® by IP injection, followed by transcardial perfusion with 10mL ice-cold PBS and 10mL 4% paraformaldehyde in PBS. Whole skulls were then isolated by decapitation and dissection and post-fixed in 4% PFA for 48hrs at 4°C. Skulls were then decalcified in 0.5M EDTA, pH 8.0 for 72hrs at 4°C, bisected at the midline, and equilibrated to 20% sucrose for 48hrs before embedding in OCT and freezing. Right hemispheres were then sectioned through the sagittal plane on a cryostat at 10µm thickness through the hippocampus and mounted directly on Xenium V1 slides. Slides were then processed and imaged on a Xenium® instrument (10X Genomics) using the stock 247-gene “Mouse Brain” panel with 84 add-on genes to identify patterns of microglial activation, pro-inflammatory cytokine genes, one-carbon metabolism, glycolysis, activation of the integrated stress response, B and T lymphocytes, and meningeal cells, as well as cell nuclei (DAPI), 18S rRNA, cell surface markers (ATP1A1, E-Cadherin, and CD45), and cytoskeletal elements (alphaSMA and Vimentin) using the Multimodal Cell Segmentation Kit (10X Genomics PN-1000661). Refinement of cell segmentation was then performed with Baysor 0.7.1 with the parameters --n-clusters=25 --min-molecules-per-cell=15 --prior-segmentation-confidence=0.5 --polygon-format=GeometryCollectionLegacy. High-level processing was then performed using Seurat 5.3.0 with SCTransform 0.4.1 (v2) normalization in R 4.4.0 using cutoffs nFeature_Xenium >= 15 & nCount_Xenium >= 20 to remove cell fragments and synapses. Images were extracted from the Xenium file bundle using Xenium Explorer 3.2.0. Cell segmentation and transcript quantification was performed using Xenium Ranger 3.1 with parameters --transcript-assignment=segmentation.csv --viz-polygons=segmentation_polygons_2d.json --units=microns. Cell annotations and IDs were exported from Seurat and imported into Xenium Explorer for identification.

For human postmortem samples, FFPE blocks from human postmortem brain tissue were sectioned at 10µm and deparaffinized in xylene followed by rehydration in 100-70% ethanol, followed by immersion in nuclease-free water. Slides were then processed and imaged on a Xenium® instrument (10X Genomics) using the stock 266-gene “Human Brain” panel with 70 add-on genes to identify patterns of microglial activation, pro-inflammatory cytokine genes, one-carbon metabolism, glycolysis, activation of the integrated stress response, B and T lymphocytes, and meningeal cells, as well as cell nuclei (DAPI), 18S rRNA, cell surface markers (ATP1A1, E-Cadherin, and CD45), and cytoskeletal elements (alphaSMA and Vimentin) using the Multimodal Cell Segmentation Kit (10X Genomics PN-1000661). Refinement of cell segmentation was then performed with Baysor 0.7.1 with the parameters --n-clusters=25 --min-molecules-per-cell=15 --prior-segmentation-confidence=0.5 --polygon-format=GeometryCollectionLegacy. High-level processing was then performed using Seurat 5.3.0 with SCTransform 0.4.2 (v2) normalization in R 4.4.0 using cutoffs nFeature_Xenium >= 10 & nCount_Xenium >= 12 to remove cell fragments and synapses. Images were extracted from the Xenium file bundle using Xenium Explorer 3.2.0. Cell segmentation and transcript quantification was performed using Xenium Ranger 3.1 with parameters --transcript-assignment=segmentation.csv --viz-polygons=segmentation_polygons_2d.json --units=microns. Cell annotations and IDs were exported from Seurat and imported into Xenium Explorer for identification.

### Spatial correlation statistics

Spatial correlation between cell types and cell states, as well as spatial autocorrelation, was determined using the spatstat ecosystem^92^, using cell centroids as determined by Xenium segmentation with Baysor refinement. Each cell state was first subsampled to 1,000 cells (or all available cells when fewer than 1,000). Two-dimensional point patterns for each cell type/state for the entire dataset were first generated using the ppp() function. Bandwidth selection for kernel density estimation was then performed using a random 10% sample of the entire dataset using the bw.ppl() function. Smoothed intensity functions were then generated for each cell type/state using the density.ppp() function using this bandwidth estimate. Spatial interactions were then determined using the inhomogeneous version of the cross K function using the Kcross.inhom() with a translation correction to handle image boundaries. A null distribution for each pair of cell types/states was obtained by randomly permuting labels across the observed cell locations (99 simulations). Statistical significance was evaluated with the global_envelope_test() function in the GET package using the extreme rank lengths method with family-wise error rate correction^93,94^. Cell type/state interaction strengths were quantified by inhomogeneous pair correlation using the pcfcross.inhom() function over a 100µm distance window. For whole-skull mouse data, only parenchymal cells were included in analysis.

### Statistical analysis and data visualization

Statistical analysis was performed using R 4.1.1^95^ with tidyverse version 2.0.0^96^ unless otherwise noted. For all comparisons, normality was first assessed using a Shapiro–Wilk test and manual examination of distributions. For parameters that exhibited a clear lack of normality, nonparametric tests were used. In cases of multiple testing, P values were corrected using FDR correction. Adjusted p-values (q-values) <0.05 were considered significant. Two-sided statistical tests were performed in all cases. Plotting was performed using ggplot 3.5.2 unless otherwise noted^97^. Comparisons for these figures were added using ggsignif 0.6.3^98^. Heatmaps were generated using ComplexHeatmap 2.15.1 with clustering using Ward’s Method (D2) with Euclidean distance as the distance metric^99^. Figure layouts were generated using patchwork 1.3.2 and edited in Adobe Illustrator 2025. In all box plots, box limits represent the interquartile range (IQR) with a center line at the median. Whiskers represent the largest point within 1.5× IQR. All points are overlaid. Outlier points are included in these overlaid points but are not shown explicitly.

### Metabolomics and LC-MS/MS

For all samples, a 5μl aliquot of the sample was used for high-resolution HPLC-tandem mass spectrometry. High-resolution HPLC-tandem mass spectrometry was performed on a Thermo Q-Exactive in line with an electrospray source and an Ultimate3000 (Thermo) series HPLC consisting of a binary pump, degasser, and auto-sampler outfitted with an Xbridge Amide column (Waters; dimensions of 4.6 mm × 100 mm and a 3.5 μm particle size). Mobile phase A contained 95% (vol/vol) water, 5% (vol/vol) acetonitrile, 10 mM ammonium hydroxide, 10 mM ammonium acetate, pH = 9.0; and mobile phase B was 100% Acetonitrile. The gradient was as follows: 0 min, 15% A; 2.5 min, 30% A; 7 min, 43% A; 16 min, 62% A; 16.1-18 min, 75% A; 18-25 min, 15% A with a flow rate of 400 μL/min. The capillary of the ESI source was set to 275 °C, with sheath gas at 45 arbitrary units, auxiliary gas at 5 arbitrary units and the spray voltage at 4.0 kV. In positive/negative polarity switching mode, an m/z scan range from 70 to 850 was chosen and MS1 data was collected at a resolution of 70,000. The automatic gain control (AGC) target was set at 1×106 and the maximum injection time was 200 ms. The top 5 precursor ions were subsequently fragmented, in a data-dependent manner, using the higher energy collisional dissociation (HCD) cell set to 30% normalized collision energy in MS2 at a resolution power of 17,500. Data acquisition and analysis were carried out by Xcalibur 4.1 software and Tracefinder 4.1 software, respectively (both from Thermo Fisher Scientific).

## Code availability

All relevant code used to produce data included in this manuscript is available on GitHub at NUPulmonary/2026_Grant (https://github.com/NUPulmonary/2026_Grant).

## Acknowledgements

We thank the many human participants and their families for their invaluable contributions to this research. This research was supported in part through a generous gift from Kimberly Querrey and Louis A. Simpson. This research was supported by the Simpson Querrey Lung Institute for Translational Science (SQLIFTS), Northwestern University. This research was also supported by the computational resources and staff contributions provided for the Quest high-performance computing facility at Northwestern University, which is jointly supported by the Office of the Provost, the Office for Research and Northwestern University Information Technology. This research was also supported in part through the computational resources and staff contributions provided by the Genomics Compute Cluster, which is jointly supported by the Feinberg School of Medicine, the Center for Genetic Medicine and Feinberg’s Department of Biochemistry and Molecular Genetics, the Office of the Provost, the Office for Research and Northwestern Information Technology. The Genomics Compute Cluster is part of Quest, Northwestern University’s high-performance computing facility, with the purpose of advancing research in genomics. We thank Jackie Milhans, Alper Kinaci, Scott Coughlin, and all members of the Research Computing and Data Services team at Northwestern for their support. Northwestern University Flow Cytometry Core Facility is supported by the National Cancer Institute Cancer Center support grant (P30 CA060553) awarded to the Robert H. Lurie Comprehensive Cancer Center. Cell sorting was performed on a BD FACS Aria SORP cell sorter purchased with the support of the National Institutes of Health (1S10OD011996-01). Integrative genomic services, including single-cell spatial transcriptomics, were performed by the Metabolomics Core Facility at Robert H. Lurie Comprehensive Cancer Center of Northwestern University. Next-generation sequencing was performed with support from the Simpson Querrey Institute for Epigenetics and the NUSeq Core Facility at Robert H. Lurie Comprehensive Cancer Center of Northwestern University.

R.A.G was supported by the Schmidt Science Fellows, in partnership with Rhodes Trust, and the Simpson Querrey Fellowship in Data Science.

T.D.G was supported by the Gefen-Querrey Brain Health Fund, The Karen Toffler Charitable Trust, and by the NIH (P30AG072977, P30AG013854).

A.V.M. was supported by the NIH (U19AI135964, P01AG049665, P01HL154998, P01HL169188, U19AI181102, R01HL153312, R01HL158139, and R01ES034350, R21AG075423), and research grants from AbbVie and Merck.

G.R.S.B. was supported by Simpson Querrey Lung Institute for Translational Science, the NIH (P01AG049665, P01HL154998, U54AG079754, R01HL147575, R01HL158139, R01HL147290, R21AG075423 and U19AI135964) and the Veterans Administration (I01CX001777).

## Supplemental Data

**Figure S1. Related to Figure 1.**
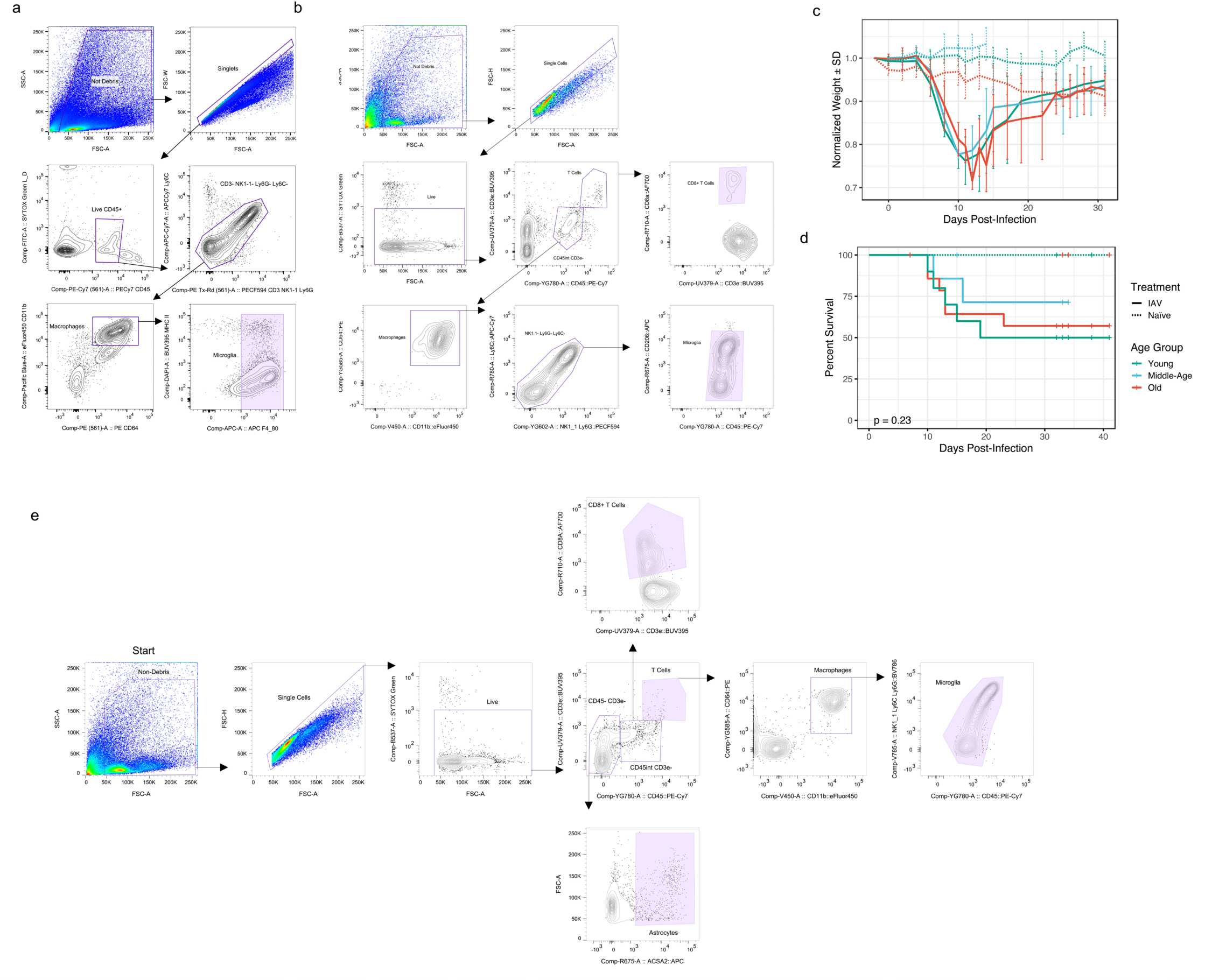
**(a)** Representative gating for flow-cytometry sorting of microglia for bulk RNA-seq experiments in the absence of zsGreen expression. **(b)** Representative gating for flow-cytometry sorting of microglia for bulk metabolomics experiments. **(c)** Body weight measurements of C57BL/6JN mice in the steady state (naïve), during acute IAV pneumonia (1 weeks post-infection), and after recovery from pneumonia (5 weeks post-infection) in young adult (4-5mo; n_naïve_ = 5, n_acute_ = 5, n_recovered_ = 5), middle-aged (12-13mo; n_naïve_ = 3, n_acute_ = 4, n_recovered_ = 5), and old mice (18-19mo; n_naïve_ = 4, n_acute_ = 8, n_recovered_ = 8). Vertices are mean weights by group as a percent of starting weight with error bars indicating standard deviation. **(d)** Kaplan-Meier survival curves for all mice in (c). No significant difference in survival was observed between IAV-treated groups (p = 0.23, Kaplan-Meier). Plus signs (+) indicate mice censored due to harvest for neuroimmune cell isolation. **(e)** Representative gating for flow-cytometry sorting of neuroimmune cells for scRNA-seq experiments.

**Figure S2. Related to Figure 2.**
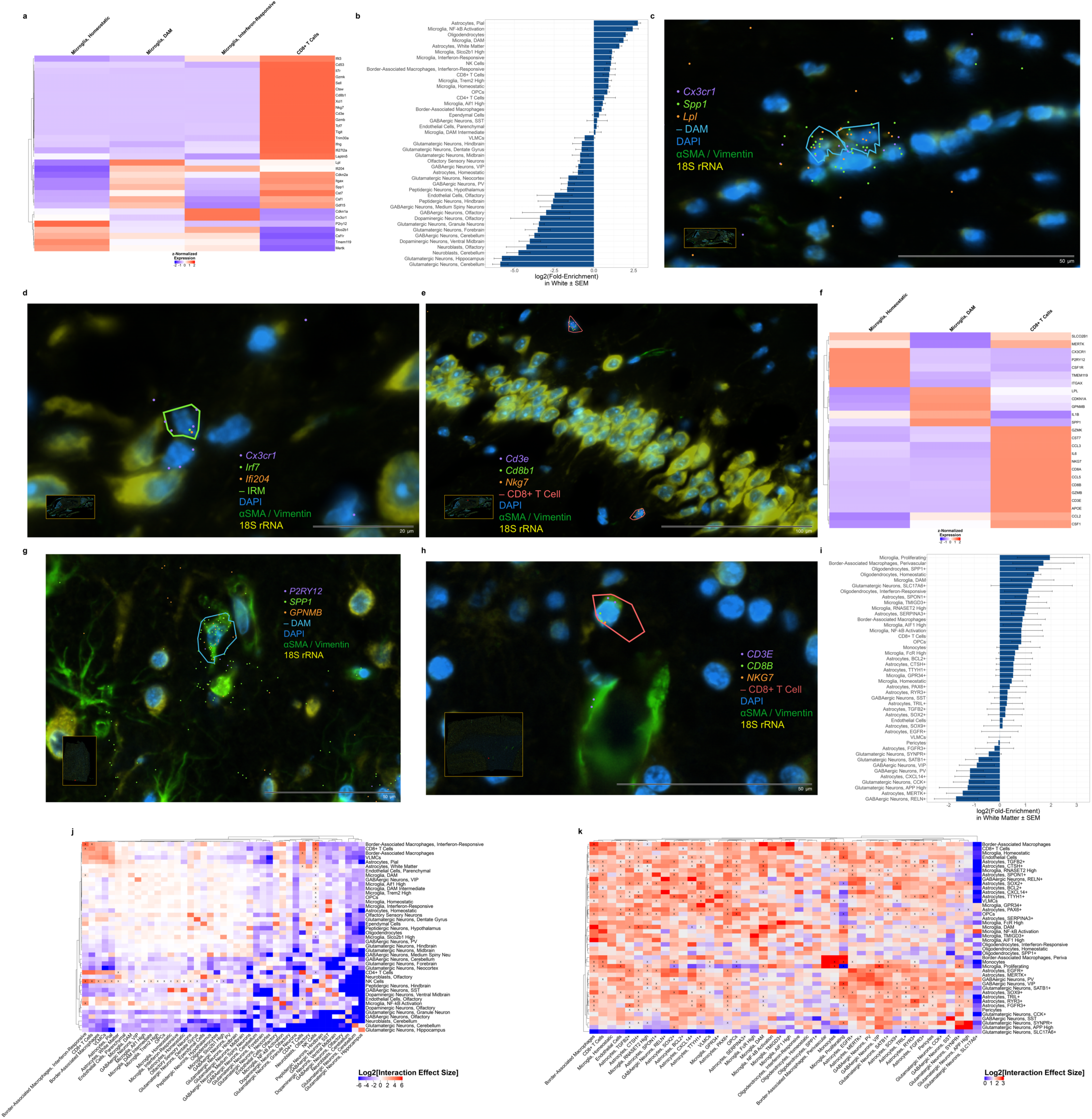
**(a)** Hierarchical clustering of mean select marker gene expression by microglia and T cell state in imaging spatial transcriptomics data from (Fig. 2a). **(b)** Log2 fold-enrichment of all detected parenchymal cell states in white matter as compared to a random distribution for each mouse in (Fig. 2a). Only parenchymal cells were included in analysis. **(c)** Representative image of DAM in imaging spatial transcriptomics data. Purple dots: *Cx3cr1*; green dots: *Spp1*; orange dots: *Lpl*; cyan polygons: DAM identified by Baysor segmentation; blue: DNA (DAPI); green: αSMA/Vimentin; yellow: 18S rRNA. **(d)** Representative image of IRM in imaging spatial transcriptomics data. Purple dots: *Cx3cr1*; green dots: *Irf7*; orange dots: *Ifi204*; green polygons: IRM identified by Baysor segmentation; blue: DNA (DAPI); green: αSMA/Vimentin; yellow: 18S rRNA. **(e)** Representative image of CD8+ T cells invading the CA2 regions of the hippocampus in an old (18mo) animal. Purple dots: *CD3e*; green dots: *Cd8b1*; orange dots: *Nkg7*; red polygons: CD8+ T cells identified by Baysor segmentation; blue: DNA (DAPI); green: αSMA/Vimentin; yellow: 18S rRNA. **(f)** Hierarchical clustering of mean select marker gene expression by microglia and T cell state from human imaging spatial transcriptomics data in (Fig. 2f). **(g)** Representative image of DAM in human imaging spatial transcriptomics data. Purple dots: *P2RY12*; green dots: *SPP1*; orange dots: *GPNMB*; cyan polygons: DAM identified by Baysor segmentation; blue: DNA (DAPI); green: αSMA/Vimentin; yellow: 18S rRNA. **(h)** Representative image of CD8+ T in human imaging spatial transcriptomics data. Purple dots: *CD3E*; green dots: *CD8B*; orange dots: *NKG7*; red polygons: CD8+ T cells identified by Baysor segmentation; blue: DNA (DAPI); green: αSMA/Vimentin; yellow: 18S rRNA. **(i)** Log2 fold-enrichment of all detected cell states in white matter as compared to a random distribution for each patient in (Fig. 2f). **(j-k)** Spatial correlation between all cell states in whole mouse skulls (j) and postmortem human MFG (j). All correlations shown are significant (q < 0.05, global envelope test against a null hypothesis of random cell state distributions with FDR correction) except those marked with an “X” (q ≥ 0.05).

**Figure S3. Related to Figure 3.**
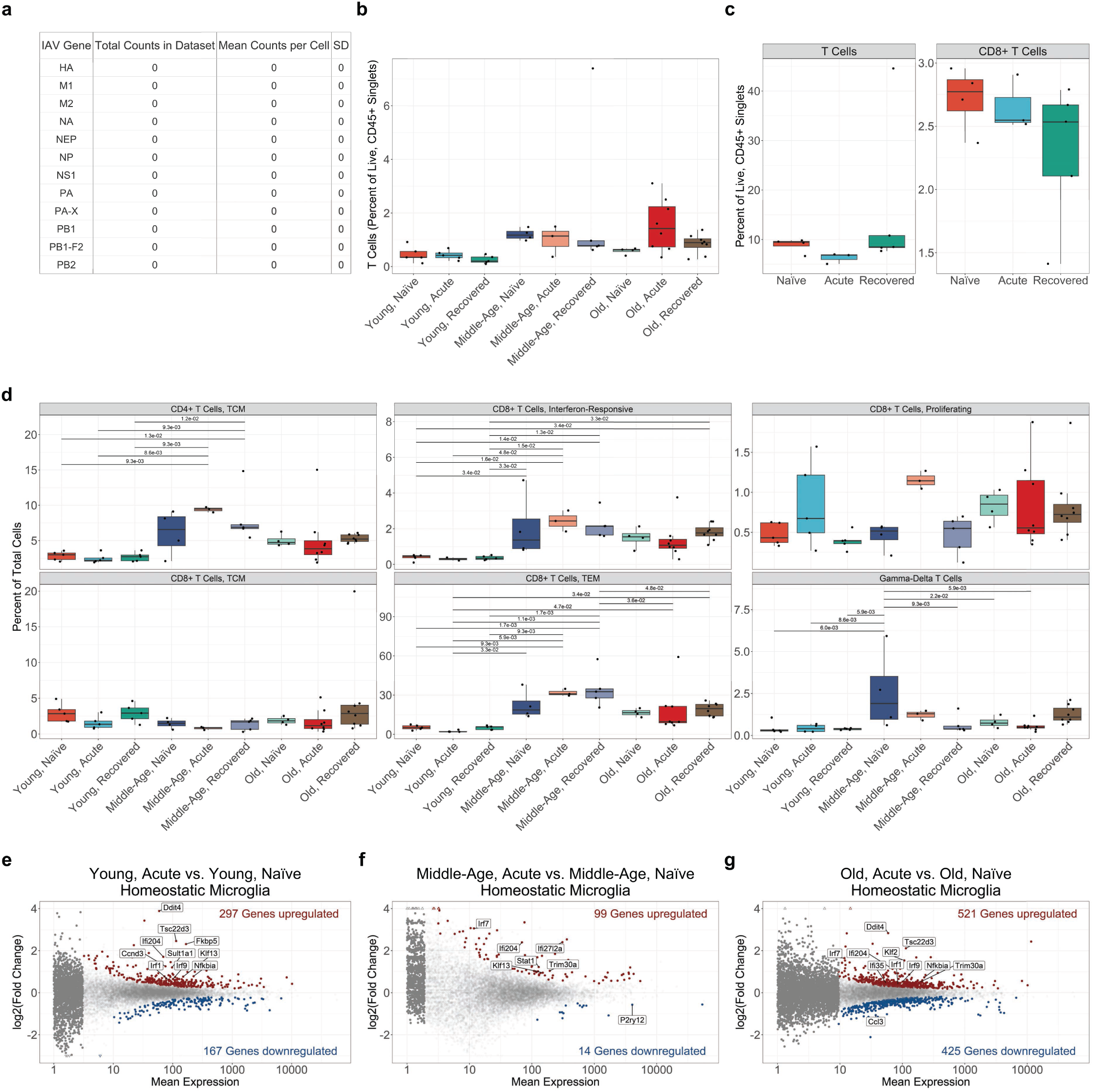
**(a)** Table of summed IAV gene expression in all cells from scRNA-seq data. **(b)** Relative flow cytometric abundances of T cells from single-cell suspensions as a proportion of live, singlet, CD45^+^ cells from whole mouse brains of C57BL/6JN mice in the steady state (naïve), during acute IAV infection (1 weeks post-infection), and after recovery from pneumonia (5 weeks post­infection) in young adult (4-5 months old; n_naïve_ = 5, n_acute_ = 5, n_recovered_ = 5), middle-aged (12-13 months old; n_naïve_ = 3, n_acute_ = 4, n_recovered_ = 5), and old age (18-19 months old; n_naïve_ = 4, n_acute_ = 8, n_recovered_ = 8). Comparisons are Wilcoxon rank sum tests with FDR correction. **(c)** Relative flow cytometric abundances of CD8+ T cells from single-cell suspensions as a proportion of live, singlet, CD45^+^ cells from whole mouse brains of C57BL/6JN mice in the steady state (naïve), during acute IAV infection (1 weeks post-infection), and after recovery from pneumonia (5 weeks post-infection) in middle-aged (12-13 months old; n_naïve_ = 3, n_acute_ = 4, n_recovered_ = 5) mice. Comparisons are Wilcoxon rank sum tests with FDR correction. **(d)** Relative abundances of selected T cell states as a percentage of total cells in scRNA-seq dataset. Comparisons are t-tests with FDR correction. **(e-g)** MA plots of pseudo-bulk RNA-seq data derived from scRNA-seq data comparing only homeostatic microglia transcriptomic responses to IAV pneumonia in (d) young adult, (e) middle-aged, and (f) old mice. Significantly (q < 0.05, Wald test) upregulated genes are shown in red, and significantly downregulated genes are shown in blue. Genes shown in gray are not significantly differentially expressed. Genes represented by triangles are outside the plot limits.

**Figure S4. Related to Figure 4.**
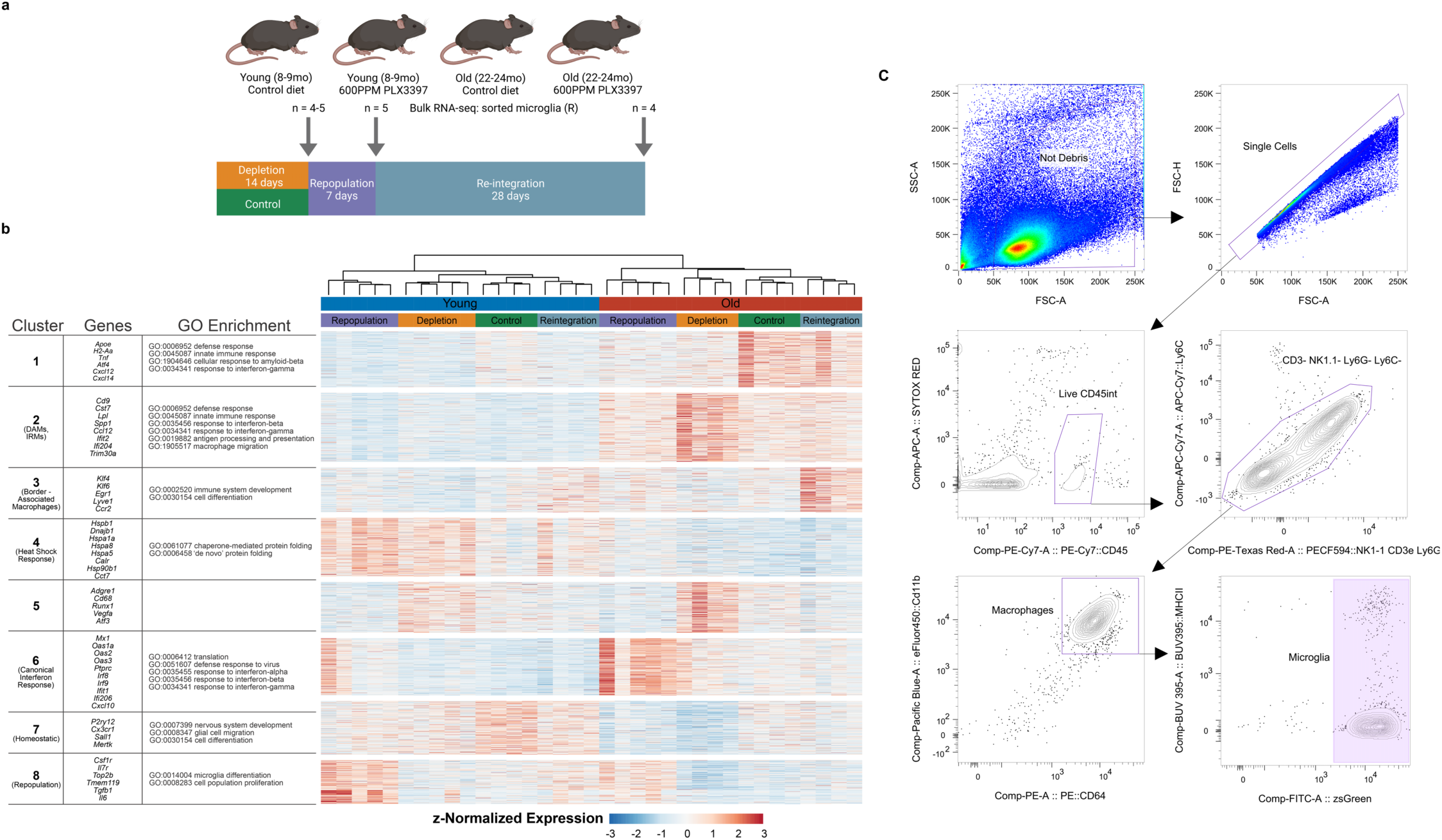
**(a)** Visual methods for microglial depletion and repopulation experiments for control chow: young adult (8mo; n = 4), old (23mo; n = 4); 14-day depletion with 600ppm PLX3397 chow: young adult (n = 4; 8mo), old (n = 4; 22mo), 7 days after PLX3397 withdrawal: young adult (n = 5; 8mo), old (n = 5; 21mo), and 30 days after PLX3397 withdrawal: young adult (n = 4; 8mo), old (n = 4; 22mo). **(b)** Hierarchical clustering of z-normalized gene expression for all mice in Figure 4a-c. All significantly variable genes (q < 0.05, likelihood ratio test) are shown. Samples (columns) were clustered using Ward’s method (D2) and genes (rows) were clustered using k-means clustering. Select genes and significantly enriched gene ontology (GO) annotations (q < 0.05, Fisher’s exact test with FDR correction) are shown at left. **(c)** Representative gating for flow cytometry sorting of microglia for bulk RNA-seq experiments in Figure 4d-e.

**Supplementary Table S1.** List of reagents used for flow cytometry sorting of mouse microglia without fluorescent reporters for bulk RNA-seq.

| Laser | Filter | Dye | Antigen | Clone | Dilution | Catalog Number | RR ID |
| --- | --- | --- | --- | --- | --- | --- | --- |
| 305 | 379/28 | BUV395 | MHC II | 2G9 | 1:500 | BD<br>743876 | AB_27418<br>27 |
| 488 | 530/30 | SYTOX<br>Green |  |  | 1:1000 | Invitrogen<br>S34860 |  |
| 405 | 450/50 | eFluor450 | CD11b | M1/70 | 1:400 | Invitrogen<br>48-0112-82 | AB_15822<br>36 |
| 640 | 670/30 | APC | F4/80 | BM8 | 1:500 | Invitrogen<br>17-4801-82 | AB_27846<br>48 |
| 640 | 780/60 | APC-Cy7 | Ly6C | HK1.4 | 1:1000 | BioLegend<br>128026 | AB_10640<br>120 |
| 552 | 575/25 | PE | CD64 | X54-5/7.1 | 1:1000 | BioLegend<br>139304 | AB_10612<br>740 |
| 552 | 610/20 | PE-CF594 | NK1.1 | PK136 | 1:2000 | BD<br>562864 | AB_27378<br>50 |
| 552 | 610/20 | PE-CF594 | CD3e | 145-2C11 | 1:333.33 | BD<br>562286 | AB_11153<br>307 |
| 552 | 610/20 | PE-CF594 | Ly6G | 1A8 | 1:2000 | BD<br>562700 | AB_27377<br>30 |
| 552 | 780/60 | PECy7 | CD45 | 30-F11 | 1:1000 | BD<br>552848 | AB_39448<br>9 |

**Supplementary Table S2.** List of reagents used for flow cytometry sorting of mouse microglia without fluorescent reporters for bulk metabolomics.

| Laser | Filter | Dye | Antigen | Clone | Dilution | Catalog<br>Number | RR ID |
| --- | --- | --- | --- | --- | --- | --- | --- |
| 305 | 379/28 | BUV395 | CD3e | 145-2C11 | 1:100 | BD<br>563565 | AB_27382<br>78 |
| 488 | 530/30 | SYTOX<br>Green |  |  | 1:1000 | Invitrogen<br>S34860 |  |
| 405 | 450/50 | eFluor450 | CD11b | M1/70 | 1:400 | Invitrogen<br>48-0112-82 | AB_15822<br>36 |
| 640 | 670/30 | APC | CD206 | C068C2 | 1:500 | BioLegend<br>141708 | AB_10900<br>231 |
| 640 | 780/60 | APC-Cy7 | Ly6C | HK1.4 | 1:1000 | BioLegend<br>128026 | AB_10640<br>120 |
| 640 | 670/30 | AF700 | CD8a | 53-6.7 | 1:333 | BD<br>557959 | AB_39695<br>9 |
| 552 | 575/25 | PE | CD64 | X54-5/7.1 | 1:1000 | BioLegend<br>139304 | AB_10612<br>740 |
| 552 | 610/20 | PE-CF594 | NK1.1 | PK136 | 1:2000 | BD<br>562864 | AB_27378<br>50 |
| 552 | 610/20 | PE-CF594 | Ly6G | 1A8 | 1:2000 | BD<br>562700 | AB_27377<br>30 |
| 552 | 780/60 | PECy7 | CD45 | 30-F11 | 1:1000 | BD<br>552848 | AB_39448<br>9 |

**Supplementary Table S3.**
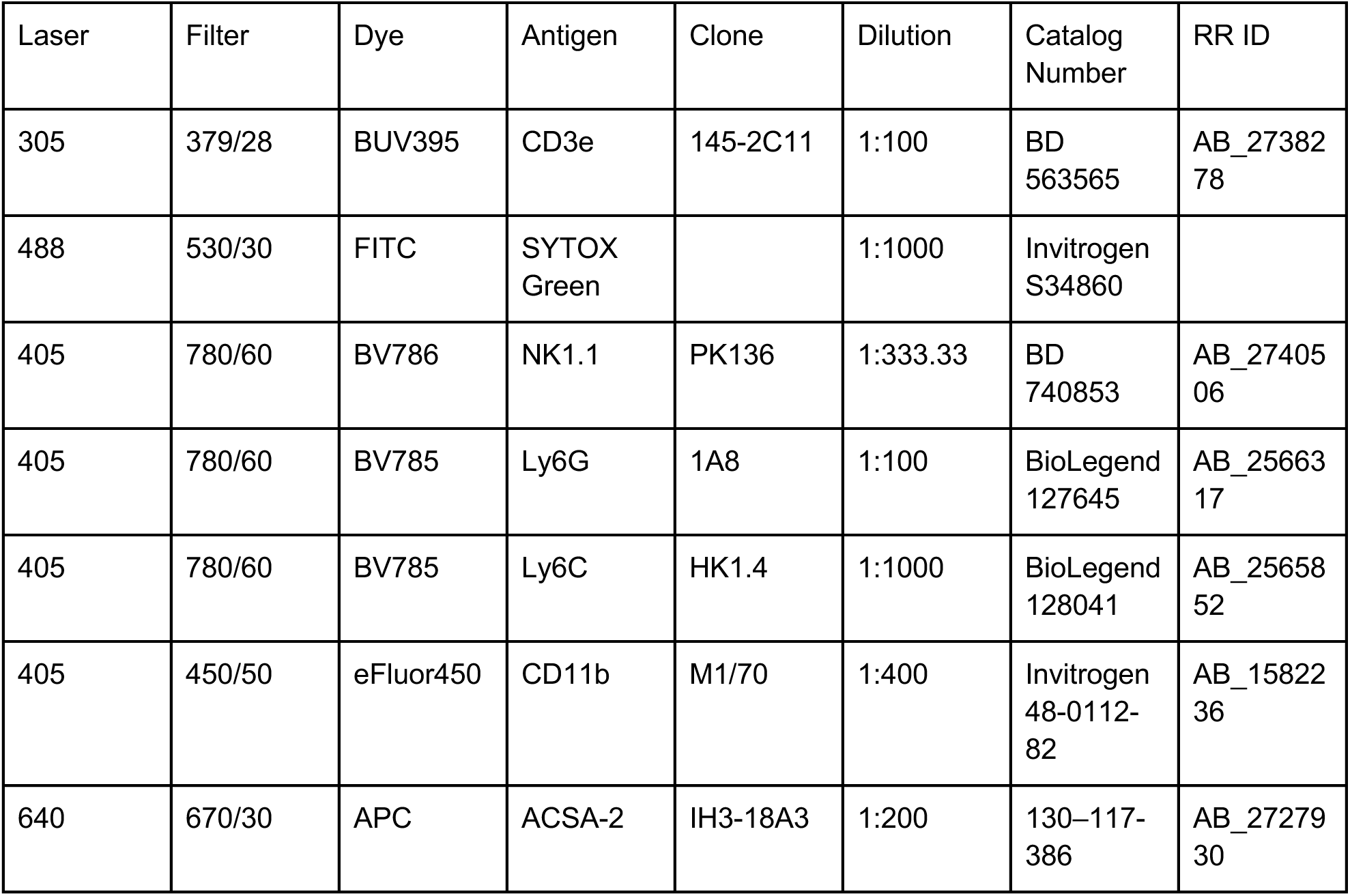
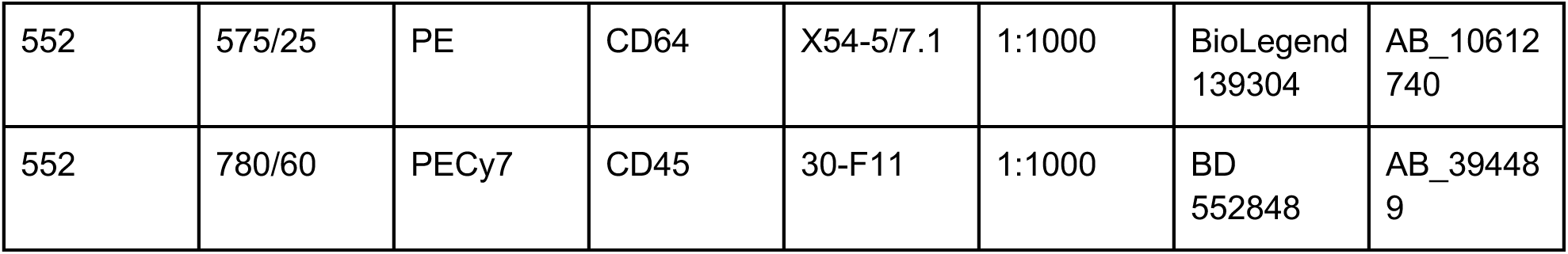
List of reagents used for flow cytometry sorting of mouse neuro-952 immune cells for scRNA-seq.

| Laser | Filter | Dye | Antigen | Clone | Dilution | Catalog Number | RR ID |
| --- | --- | --- | --- | --- | --- | --- | --- |
| 305 | 379/28 | BUV395 | CD3e | 145-2C11 | 1:100 | BD<br>563565 | AB_27382<br>78 |
| 488 | 530/30 | FITC | SYTOX<br>Green |  | 1:1000 | Invitrogen<br>S34860 |  |
| 405 | 780/60 | BV786 | NK1.1 | PK136 | 1:333.33 | BD<br>740853 | AB_27405<br>06 |
| 405 | 780/60 | BV785 | Ly6G | 1A8 | 1:100 | BioLegend<br>127645 | AB_25663<br>17 |
| 405 | 780/60 | BV785 | Ly6C | HK1.4 | 1:1000 | BioLegend<br>128041 | AB_25658<br>52 |
| 405 | 450/50 | eFluor450 | CD11b | M1/70 | 1:400 | Invitrogen<br>48-0112-82 | AB_15822<br>36 |
| 640 | 670/30 | APC | ACSA-2 | IH3-18A3 | 1:200 | 130-117-386 | AB_27279<br>30 |
| 552 | 575/25 | PE | CD64 | X54-5/7.1 | 1:1000 | BioLegend<br>139304 | AB_10612<br>740 |
| 552 | 780/60 | PECy7 | CD45 | 30-F11 | 1:1000 | BD<br>552848 | AB_39448<br>9 |

**Supplementary Table S4.**
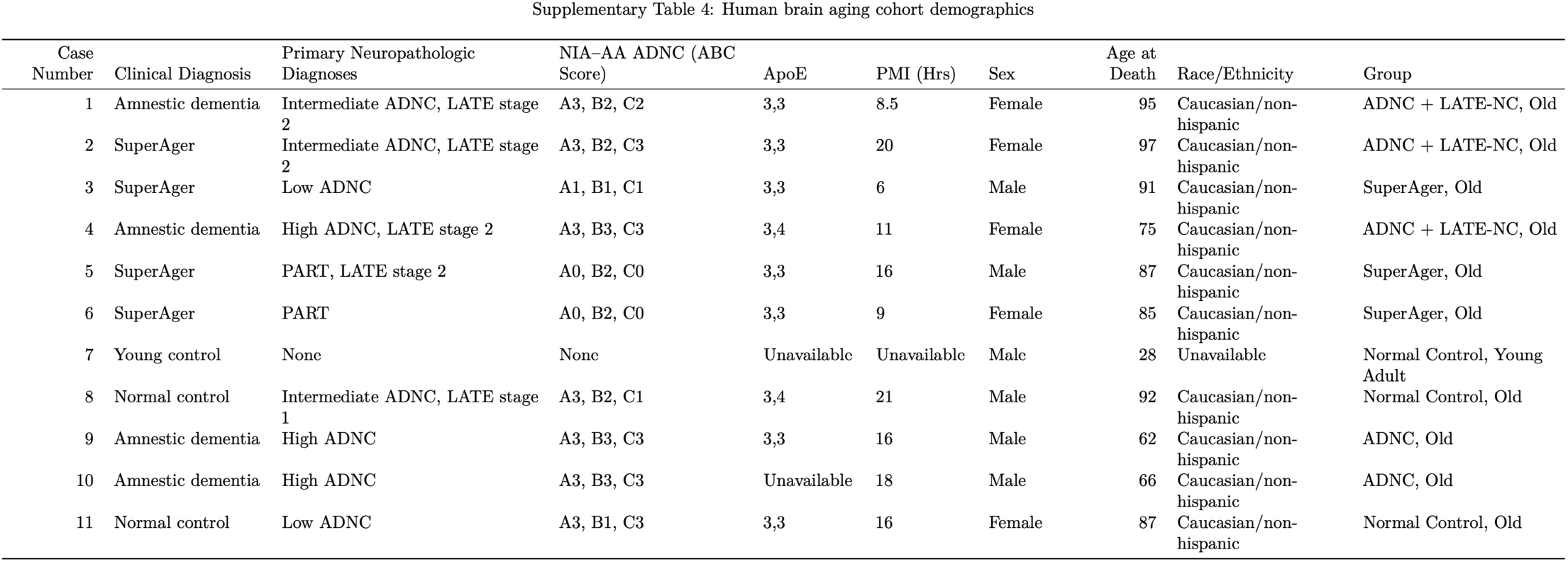
Cohort demographics for human MFG spatial transcriptomics analysis.

**Supplementary Table S5.**
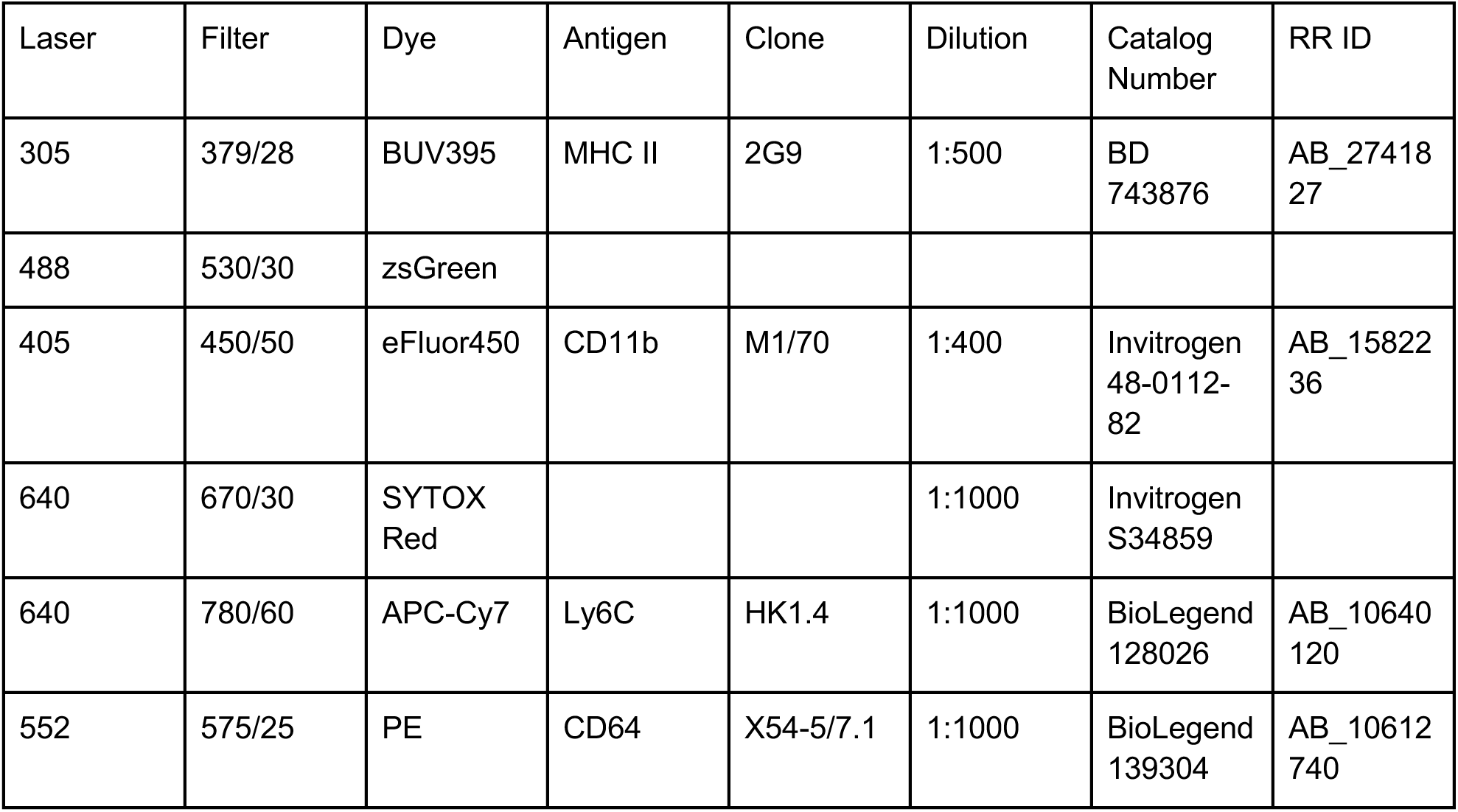
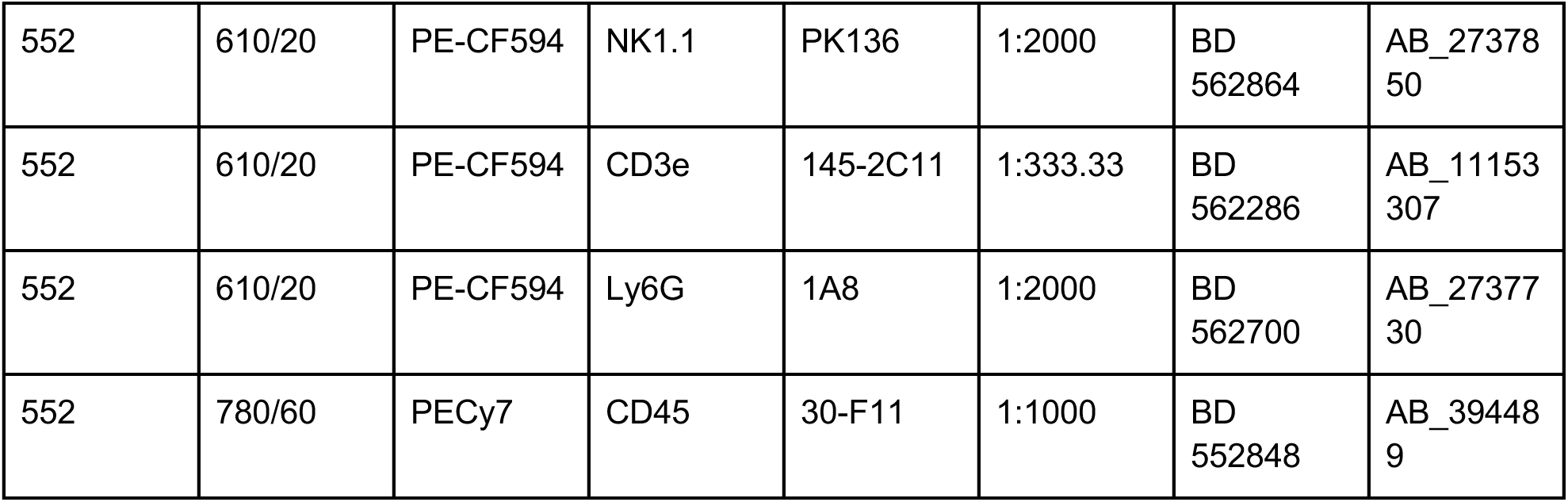
List of reagents used for flow cytometry sorting of mouse microglia expressing zsGreen for bulk RNA-seq.

| Laser | Filter | Dye | Antigen | Clone | Dilution | Catalog Number | RR ID |
| --- | --- | --- | --- | --- | --- | --- | --- |
| 305 | 379/28 | BUV395 | MHC II | 2G9 | 1:500 | BD<br>743876 | AB_27418<br>27 |
| 488 | 530/30 | zsGreen |  |  |  |  |  |
| 405 | 450/50 | eFluor450 | CD11b | M1/70 | 1:400 | Invitrogen<br>48-0112-82 | AB_15822<br>36 |
| 640 | 670/30 | SYTOX Red |  |  | 1:1000 | Invitrogen<br>S34859 |  |
| 640 | 780/60 | APC-Cy7 | Ly6C | HK1.4 | 1:1000 | BioLegend<br>128026 | AB_10640<br>120 |
| 552 | 575/25 | PE | CD64 | X54-5/7.1 | 1:1000 | BioLegend<br>139304 | AB_10612<br>740 |
| 552 | 610/20 | PE-CF594 | NK1.1 | PK136 | 1:2000 | BD<br>562864 | AB_27378<br>50 |
| 552 | 610/20 | PE-CF594 | CD3e | 145-2C11 | 1:333.33 | BD<br>562286 | AB_11153<br>307 |
| 552 | 610/20 | PE-CF594 | Ly6G | 1A8 | 1:2000 | BD<br>562700 | AB_27377<br>30 |
| 552 | 780/60 | PECy7 | CD45 | 30-F11 | 1:1000 | BD<br>552848 | AB_39448<br>9 |

## References

1. Shah, F. A. et al. Bidirectional Relationship between Cognitive Function and Pneumonia. Am J Respir Crit Care Med 188, 586–592 (2013).

2. Dunn, N., Mullee, M., Perry, V. H. & Holmes, C. Association between Dementia and Infectious Disease: Evidence from a Case-Control Study. Alzheimer Disease & Associated Disorders 19, 91–94 (2005).

3. Davydow, D. S., Hough, C. L., Levine, D. A., Langa, K. M. & Iwashyna, T. J. Functional Disability, Cognitive Impairment, and Depression Following Hospitalization for Pneumonia. Am J Med 126, 615–624.e5 (2013).

4. Wang, W., Tang, J. & Wei, F. Updated understanding of the outbreak of 2019 novel coronavirus (2019-nCoV) in Wuhan, China. Journal of Medical Virology 92, 441–447 (2020).

5. Khan, S. & Gomes, J. Neuropathogenesis of SARS-CoV-2 infection. eLife 9, e59136 (2020).

6. Koralnik, I. J. & Tyler, K. L. COVID-19: A Global Threat to the Nervous System. Annals of Neurology 88, 1–11 (2020).

7. Levine, K. S. et al. Virus exposure and neurodegenerative disease risk across national biobanks. Neuron 111, 1086–1093.e2 (2023).

8. Spudich, S. & Nath, A. Nervous system consequences of COVID-19. Science 10.1126/science.abm2052 (2022) doi:10.1126/science.abm2052.

9. Kuiken, T. & Taubenberger, J. K. Pathology of human influenza revisited. Vaccine 26, D59–D66 (2008).

10. Budinger, G. R. S., Misharin, A. V., Ridge, K. M., Singer, B. D. & Wunderink, R. G. Distinctive features of severe SARS-CoV-2 pneumonia. Journal of Clinical Investigation 131, e149412 (2021).

11. Jurgens, H. A., Amancherla, K. & Johnson, R. W. Influenza Infection Induces Neuroinflammation, Alters Hippocampal Neuron Morphology, and Impairs Cognition in Adult Mice. J. Neurosci. 32, 3958–3968 (2012).

12. Dinnon, K. H. et al. A mouse-adapted model of SARS-CoV-2 to test COVID-19 countermeasures. Nature 586, 560–566 (2020).

13. Fernández-Castañeda, A. et al. Mild respiratory COVID can cause multi-lineage neural cell and myelin dysregulation. Cell 10.1016/j.cell.2022.06.008 (2022) doi:10.1016/j.cell.2022.06.008.

14. Sadasivan, S., Zanin, M., O’Brien, K., Schultz-Cherry, S. & Smeyne, R. J. Induction of Microglia Activation after Infection with the Non-Neurotropic A/CA/04/2009 H1N1 Influenza Virus. PLoS One 10, (2015).

15. Hosseini, S. et al. Long-Term Neuroinflammation Induced by Influenza A Virus Infection and the Impact on Hippocampal Neuron Morphology and Function. J. Neurosci. 38, 3060–3080 (2018).

16. Grant, R. A., et al. Prolonged exposure to lung-derived cytokines is associated with activation of microglia in patients with COVID-19. JCI Insight 9, (2024).

17. Schwabenland, M. et al. Deep spatial profiling of human COVID-19 brains reveals neuroinflammation with distinct microanatomical microglia-T-cell interactions. Immunity 54, 1594–1610.e11 (2021).

18. Fullard, J. F. et al. Single-nucleus transcriptome analysis of human brain immune response in patients with severe COVID-19. Genome Med 13, 118 (2021).

19. Lee, M. H. et al. Neurovascular injury with complement activation and inflammation in COVID-19. Brain awac151 (2022) doi:10.1093/brain/awac151.

20. Cosentino, G. et al. Neuropathological findings from COVID-19 patients with neurological symptoms argue against a direct brain invasion of SARS-CoV-2: A critical systematic review. European Journal of Neurology 28, 3856–3865 (2021).

21. Soung, A. L. et al. COVID-19 induces CNS cytokine expression and loss of hippocampal neurogenesis. Brain 145, 4193–4201 (2022).

22. Keren-Shaul, H. et al. A Unique Microglia Type Associated with Restricting Development of Alzheimer’s Disease. Cell 169, 1276–1290.e17 (2017).

23. Deczkowska, A. et al. Disease-Associated Microglia: A Universal Immune Sensor of Neurodegeneration. Cell 173, 1073–1081 (2018).

24. Kaya, T. et al. CD8+ T cells induce interferon-responsive oligodendrocytes and microglia in white matter aging. Nat Neurosci 1–12 (2022) doi:10.1038/s41593-022-01183-6.

25. Safaiyan, S. et al. White matter aging drives microglial diversity. Neuron 109, 1100–1117.e10 (2021).

26. Hahn, O. et al. Atlas of the aging mouse brain reveals white matter as vulnerable foci. Cell 186, 4117–4133.e22 (2023).

27. Gefen, T. et al. Activated Microglia in Cortical White Matter Across Cognitive Aging Trajectories. Front Aging Neurosci 11, 94 (2019).

28. O’Neil, S. M., Witcher, K. G., McKim, D. B. & Godbout, J. P. Forced turnover of aged microglia induces an intermediate phenotype but does not rebalance CNS environmental cues driving priming to immune challenge. Acta Neuropathologica Communications 6, 129 (2018).

29. Henry, C. J., Huang, Y., Wynne, A. M. & Godbout, J. P. Peripheral lipopolysaccharide (LPS) challenge promotes microglial hyperactivity in aged mice that is associated with exaggerated induction of both pro-inflammatory IL-1β and anti-inflammatory IL-10 cytokines. Brain, Behavior, and Immunity 23, 309–317 (2009).

30. Perry, V. H., Cunningham, C. & Holmes, C. Systemic infections and inflammation affect chronic neurodegeneration. Nat Rev Immunol 7, 161–167 (2007).

31. Elmore, M. R. P. et al. Replacement of microglia in the aged brain reverses cognitive, synaptic, and neuronal deficits in mice. Aging Cell 17, e12832 (2018).

32. Wendeln, A.-C. et al. Innate immune memory in the brain shapes neurological disease hallmarks. Nature 556, 332–338 (2018).

33. Cunningham, C., Wilcockson, D. C., Campion, S., Lunnon, K. & Perry, V. H. Central and Systemic Endotoxin Challenges Exacerbate the Local Inflammatory Response and Increase Neuronal Death during Chronic Neurodegeneration. J. Neurosci. 25, 9275–9284 (2005).

34. Aged C57BL/6J Mice for Research Studies. https://resources.jax.org/white-papers/whitepaper-aged-b6 (2023).

35. Hammond, T. R. et al. Single-Cell RNA Sequencing of Microglia throughout the Mouse Lifespan and in the Injured Brain Reveals Complex Cell-State Changes. Immunity 50, 253–271.e6 (2019).

36. Mao, Y., Shi, D., Li, G. & Jiang, P. Citrulline depletion by ASS1 is required for proinflammatory macrophage activation and immune responses. Molecular Cell 82, 527–541.e7 (2022).

37. Suryadevara, V. et al. SenNet recommendations for detecting senescent cells in different tissues. Nat Rev Mol Cell Biol 25, 1001–1023 (2024).

38. Saul, D. et al. A new gene set identifies senescent cells and predicts senescence-associated pathways across tissues. Nat Commun 13, 4827 (2022).

39. Liberzon, A. et al. Molecular signatures database (MSigDB) 3.0. Bioinformatics 27, 1739–1740 (2011).

40. Lampropoulou, V. et al. Itaconate Links Inhibition of Succinate Dehydrogenase with Macrophage Metabolic Remodeling and Regulation of Inflammation. Cell Metabolism 24, 158–166 (2016).

41. Strelko, C. L. et al. Itaconic Acid Is a Mammalian Metabolite Induced during Macrophage Activation. J. Am. Chem. Soc. 133, 16386–16389 (2011).

42. Cordes, T. et al. Immunoresponsive Gene 1 and Itaconate Inhibit Succinate Dehydrogenase to Modulate Intracellular Succinate Levels. Journal of Biological Chemistry 291, 14274–14284 (2016).

43. O’Neill, L. A. J. & Artyomov, M. N. Itaconate: the poster child of metabolic reprogramming in macrophage function. Nat Rev Immunol 19, 273–281 (2019).

44. Chandel, N. S. Navigating Metabolism. vol. 517 (Cold Spring Harbor Laboratory Press Cold Spring Harbor, New York, 2015).

45. McQuattie-Pimentel, A. C. et al. The lung microenvironment shapes a dysfunctional response of alveolar macrophages in aging. J Clin Invest 131, (2021).

46. Byrd-Leotis, L. et al. Influenza binds phosphorylated glycans from human lung. Science Advances 5, eaav2554 (2019).

47. Radigan, K. A., Misharin, A. V., Chi, M. & Budinger, G. S. Modeling human influenza infection in the laboratory. Infect Drug Resist 8, 311–320 (2015).

48. Runyan, C. E. et al. Impaired phagocytic function in CX3CR1+ tissue-resident skeletal muscle macrophages prevents muscle recovery after influenza A virus-induced pneumonia in old mice. Aging Cell 19, e13180 (2020).

49. Dolan, M.-J. et al. Spatiotemporal Analysis of Remyelination Reveals a Concerted Interferon-Responsive Glial State That Coordinates Immune Infiltration. 2025.04.22.649486 Preprint at 10.1101/2025.04.22.649486 (2025).

50. Harrison, T. M., Weintraub, S., Mesulam, M.-M. & Rogalski, E. Superior Memory and Higher Cortical Volumes in Unusually Successful Cognitive Aging. Journal of the International Neuropsychological Society 18, 1081–1085 (2012).

51. Hyman, B. T. et al. National Institute on Aging–Alzheimer’s Association guidelines for the neuropathologic assessment of Alzheimer’s disease. Alzheimer’s & dementia 8, 1–13 (2012).

52. Montine, T. J. et al. National Institute on Aging–Alzheimer’s Association guidelines for the neuropathologic assessment of Alzheimer’s disease: a practical approach. Acta neuropathologica 123, 1–11 (2012).

53. McKhann, G. M. et al. The diagnosis of dementia due to Alzheimer’s disease: recommendations from the National Institute on Aging-Alzheimer’s Association workgroups on diagnostic guidelines for Alzheimer’s disease. Alzheimer’s & dementia 7, 263–269 (2011).

54. Josephs, K. A. et al. Updated TDP-43 in Alzheimer’s disease staging scheme. Acta neuropathologica 131, 571–585 (2016).

55. Nelson, P. T. et al. Limbic-predominant age-related TDP-43 encephalopathy (LATE): consensus working group report. Brain 142, 1503–1527 (2019).

56. Ayroldi, E. & Riccardi, C. Glucocorticoid-induced leucine zipper (GILZ): a new important mediator of glucocorticoid action. The FASEB Journal 23, 3649–3658 (2009).

57. Zannas, A. S., Wiechmann, T., Gassen, N. C. & Binder, E. B. Gene–Stress–Epigenetic Regulation of FKBP5: Clinical and Translational Implications. Neuropsychopharmacol 41, 261–274 (2016).

58. Wang, J.-C., Gray, N. E., Kuo, T. & Harris, C. A. Regulation of triglyceride metabolism by glucocorticoid receptor. Cell Biosci 2, 19 (2012).

59. Polman, J. A. E. et al. Glucocorticoids Modulate the mTOR Pathway in the Hippocampus: Differential Effects Depending on Stress History. Endocrinology 153, 4317–4327 (2012).

60. Ip, W. K. E., Hoshi, N., Shouval, D. S., Snapper, S. & Medzhitov, R. Anti-inflammatory effect of IL-10 mediated by metabolic reprogramming of macrophages. Science 356, 513–519 (2017).

61. Godot, V. et al. Dexamethasone and IL-10 stimulate glucocorticoid-induced leucine zipper synthesis by human mast cells. Allergy 61, 886–890 (2006).

62. Elmore, M. R. P. et al. Colony-Stimulating Factor 1 Receptor Signaling Is Necessary for Microglia Viability, Unmasking a Microglia Progenitor Cell in the Adult Brain. Neuron 82, 380–397 (2014).

63. Liu, M. et al. GPNMB and ATP6V1A interact to mediate microglia phagocytosis of multiple types of pathological particles. Cell Reports 44, 115343 (2025).

64. Saade, M., Araujo de Souza, G., Scavone, C. & Kinoshita, P. F. The Role of GPNMB in Inflammation. Front. Immunol. 12, (2021).

65. Zhang, S. et al. Efferocytosis Fuels Requirements of Fatty Acid Oxidation and the Electron Transport Chain to Polarize Macrophages for Tissue Repair. Cell Metabolism 29, 443–456.e5 (2019).

66. Cai, B. et al. MerTK cleavage limits proresolving mediator biosynthesis and exacerbates tissue inflammation. PNAS 113, 6526–6531 (2016).

67. Huang, Y. & Lemke, G. Early death in a mouse model of Alzheimer’s disease exacerbated by microglial loss of TAM receptor signaling. Proceedings of the National Academy of Sciences 119, e2204306119 (2022).

68. Dolan, M.-J. et al. Exposure of iPSC-derived human microglia to brain substrates enables the generation and manipulation of diverse transcriptional states in vitro. Nat Immunol 24, 1382–1390 (2023).

69. Anderson, S. R. et al. Neuronal apoptosis drives remodeling states of microglia and shifts in survival pathway dependence. eLife 11, e76564 (2022).

70. Runyan, C. E. et al. Tissue-resident skeletal muscle macrophages promote recovery from viral pneumonia-induced sarcopenia in normal aging. 2025.01.09.631996 Preprint at 10.1101/2025.01.09.631996 (2025).

71. Goldmann, T. et al. Origin, fate and dynamics of macrophages at central nervous system interfaces. Nat Immunol 17, 797–805 (2016).

72. Alliot, F., Godin, I. & Pessac, B. Microglia derive from progenitors, originating from the yolk sac, and which proliferate in the brain. Developmental Brain Research 117, 145–152 (1999).

73. Raj, D. D. A. et al. Priming of microglia in a DNA-repair deficient model of accelerated aging. Neurobiology of Aging 35, 2147–2160 (2014).

74. Gal, A. et al. Mutations in MERTK, the human orthologue of the RCS rat retinal dystrophy gene, cause retinitis pigmentosa. Nat Genet 26, 270–271 (2000).

75. Chung, W.-S. et al. Astrocytes mediate synapse elimination through MEGF10 and MERTK pathways. Nature 504, 394–400 (2013).

76. Saito, K. et al. High-Resolution Spatial Profiling of Microglia Reveals Proximity Associated Immunometabolic Reprogramming in Alzheimer’s Disease. 2025.05.16.654329 Preprint at 10.1101/2025.05.16.654329 (2025).

77. Pálovics, R. et al. Molecular hallmarks of heterochronic parabiosis at single-cell resolution. Nature 603, 309–314 (2022).

78. Yona, S. et al. Fate Mapping Reveals Origins and Dynamics of Monocytes and Tissue Macrophages under Homeostasis. Immunity 38, 79–91 (2013).

79. Madisen, L. et al. A robust and high-throughput Cre reporting and characterization system for the whole mouse brain. Nat Neurosci 13, 133–140 (2010).

80. Li, Y. et al. The role of endothelial MERTK during the inflammatory response in lungs. PLoS One 14, e0225051 (2019).

81. Coates, B. M. et al. Inflammatory Monocytes Drive Influenza A Virus–Mediated Lung Injury in Juvenile Mice. The Journal of Immunology 200, 2391–2404 (2018).

82. Morales-Nebreda, L. et al. Intratracheal administration of influenza virus is superior to intranasal administration as a model of acute lung injury. Journal of Virological Methods 209, 116–120 (2014).

83. Love, M. I., Huber, W. & Anders, S. Moderated estimation of fold change and dispersion for RNA-seq data with DESeq2. Genome Biology 15, 550 (2014).

84. Wolock, S. L., Lopez, R. & Klein, A. M. Scrublet: Computational Identification of Cell Doublets in Single-Cell Transcriptomic Data. Cell Systems 8, 281–291.e9 (2019).

85. Satija, R., Farrell, J. A., Gennert, D., Schier, A. F. & Regev, A. Spatial reconstruction of single-cell gene expression data. Nature Biotechnology 33, 495–502 (2015).

86. Lopez, R., Regier, J., Cole, M. B., Jordan, M. I. & Yosef, N. Deep generative modeling for single-cell transcriptomics. Nat Methods 15, 1053–1058 (2018).

87. Mangiola, S., Doyle, M. A. & Papenfuss, A. T. Interfacing Seurat with the R tidy universe. Bioinformatics 37, 4100–4107 (2021).

88. Hafemeister, C. & Satija, R. Normalization and Variance Stabilization of Single-Cell RNA-Seq Data Using Regularized Negative Binomial Regression. http://biorxiv.org/lookup/doi/10.1101/576827 (2019) doi:10.1101/576827.

89. Traag, V. A., Waltman, L. & van Eck, N. J. From Louvain to Leiden: guaranteeing well-connected communities. Sci Rep 9, 5233 (2019).

90. Schmidt, M. Rey Auditory Verbal Learning Test: RAVLT : A Handbook. (Western Psychological Services, 1996).

91. Gefen, T. et al. Morphometric and Histologic Substrates of Cingulate Integrity in Elders with Exceptional Memory Capacity. J. Neurosci. 35, 1781–1791 (2015).

92. Baddeley, A. & Turner, R. spatstat: An R Package for Analyzing Spatial Point Patterns. Journal of Statistical Software 12, 1–42 (2005).

93. Myllymäki, M., Mrkvička, T., Grabarnik, P., Seijo, H. & Hahn, U. Global Envelope Tests for Spatial Processes. J. R. Stat. Soc. Ser. B. Stat. Methodol. 79, 381–404 (2017).

94. Myllymäki, M. & Mrkvička, T. GET: Global Envelopes in R. Journal of Statistical Software 111, 1–40 (2024).

95. Ihaka, R. & Gentleman, R. R: A Language for Data Analysis and Graphics. Journal of Computational and Graphical Statistics 5, 299–314 (1996).

96. Wickham, H. et al. Welcome to the Tidyverse. Journal of Open Source Software 4, 1686 (2019).

97. Wickham, H. Ggplot2: Elegant Graphics for Data Analysis. (Springer, 2016).

98. Constantin, A. & Patil, I. ggsignif: R package for displaying significance brackets for “ggplot2.” PsyArxiv (2021).

99. Gu, Z., Eils, R. & Schlesner, M. Complex heatmaps reveal patterns and correlations in multidimensional genomic data. Bioinformatics 32, 2847–2849 (2016).

