## Supplementary figures and images for "Influenza A infection accelerates disease-associated microglia formation during physiological aging"

### Figure 1

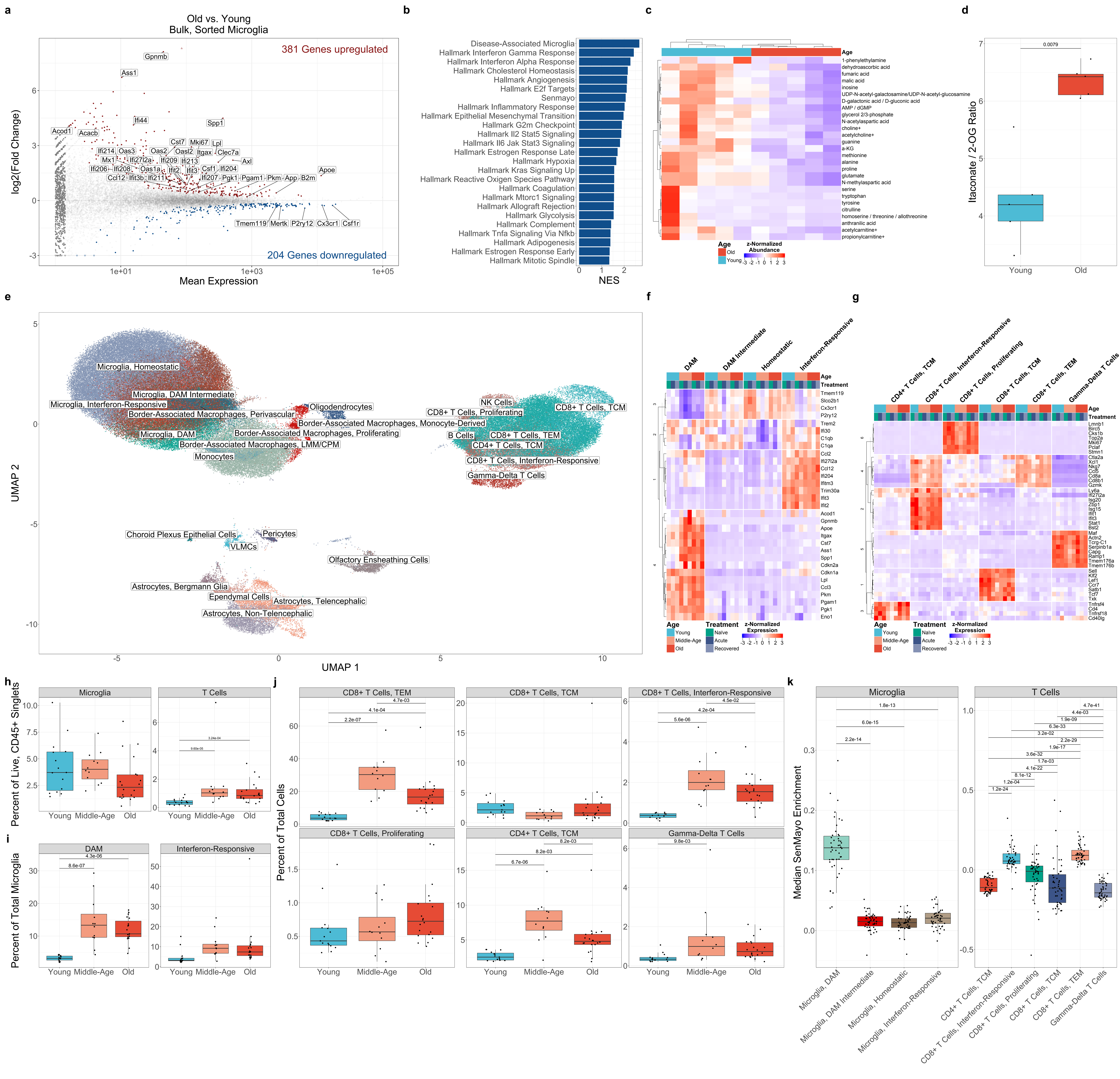

### Figure 2

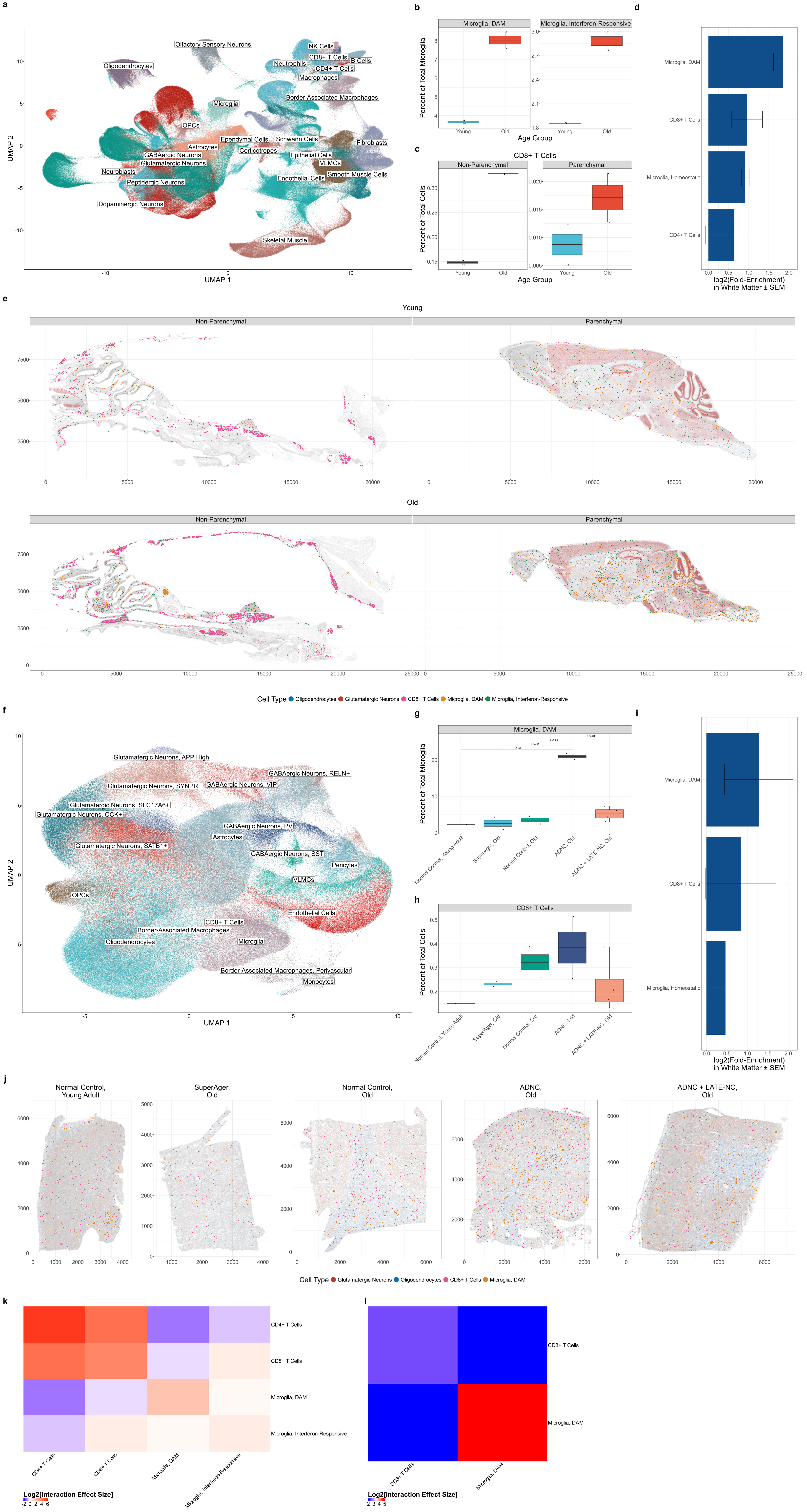

### Figure 3

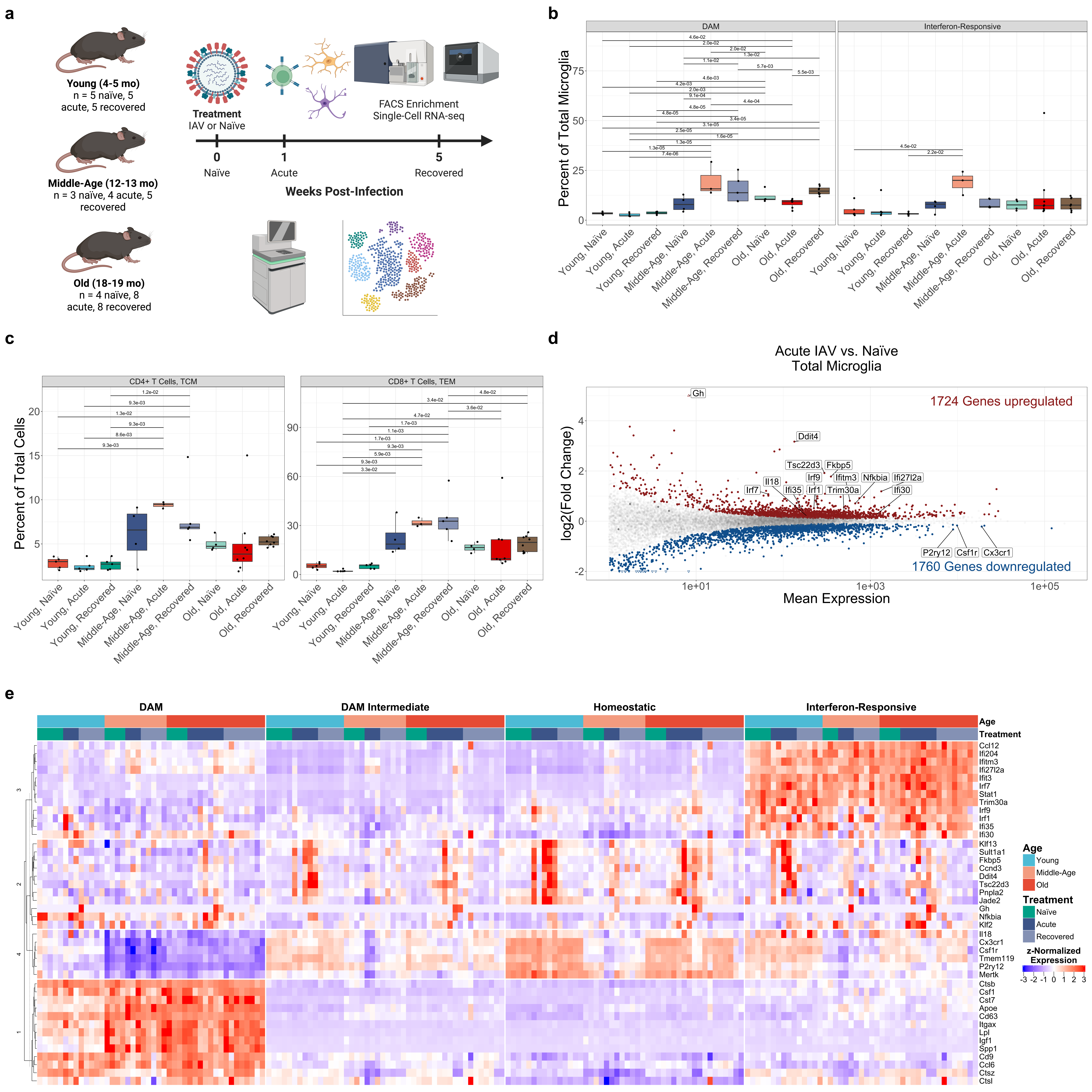

### Figure 4

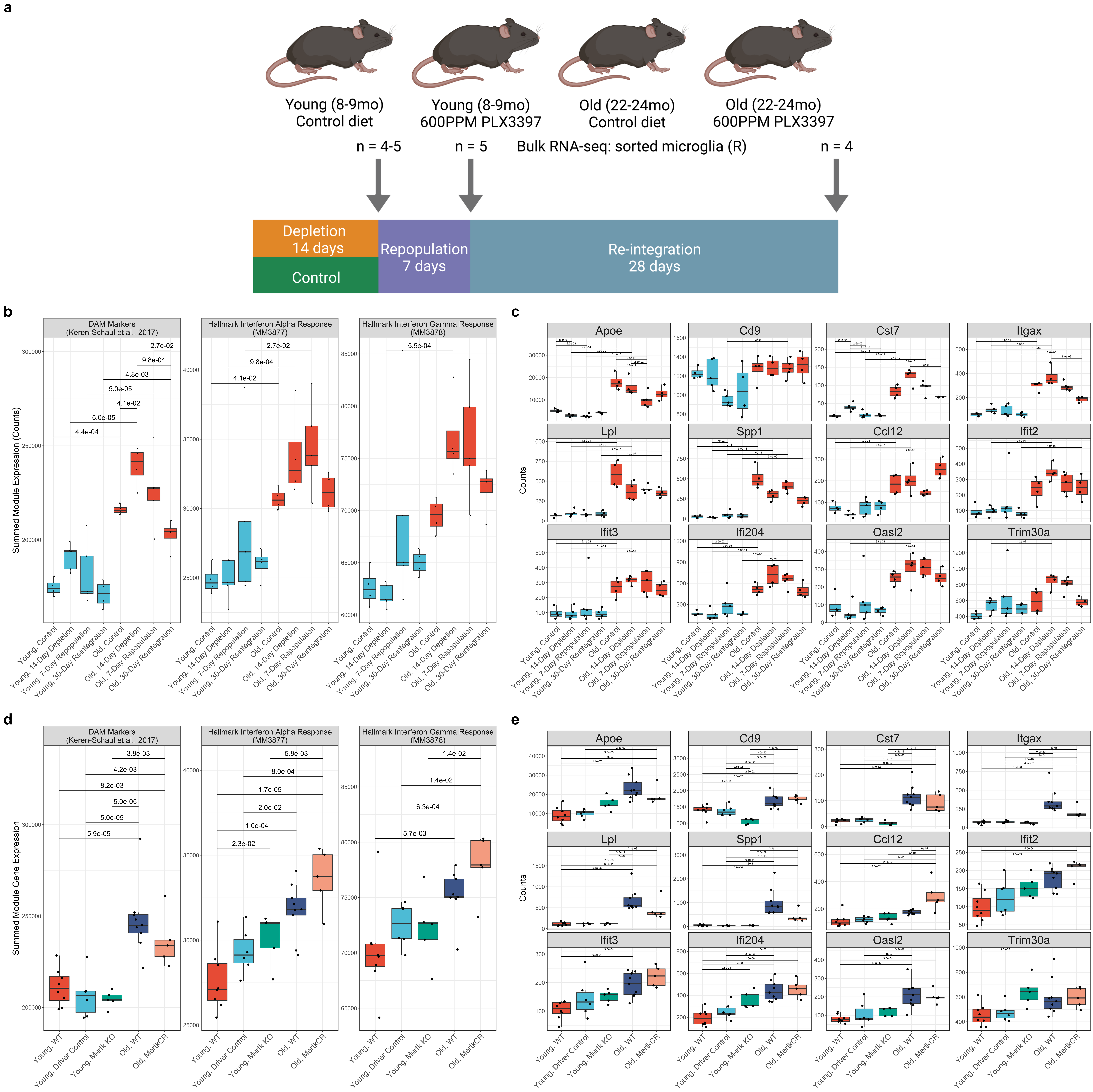

### Supplementary Figure S1

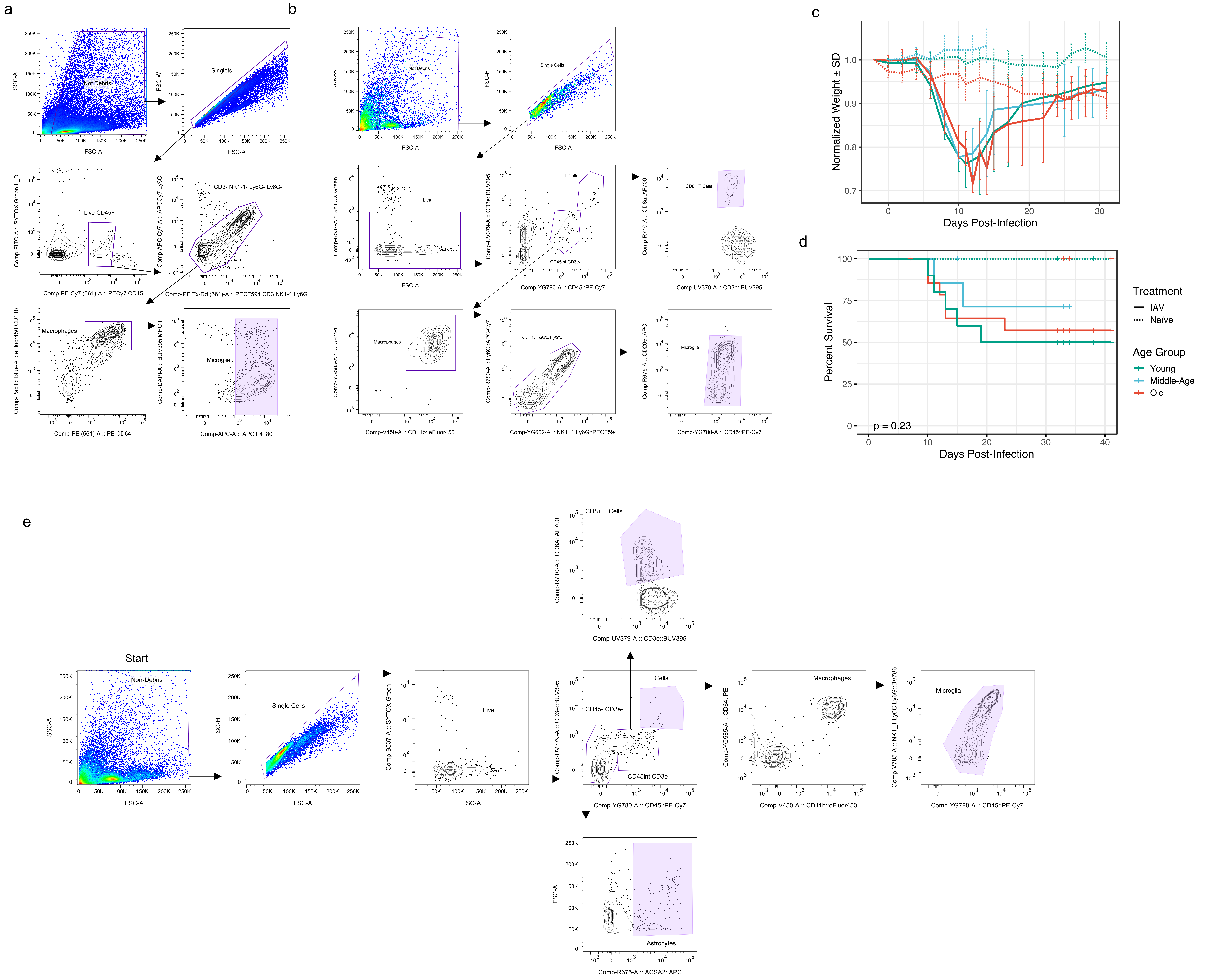

### Supplementary Figure S2

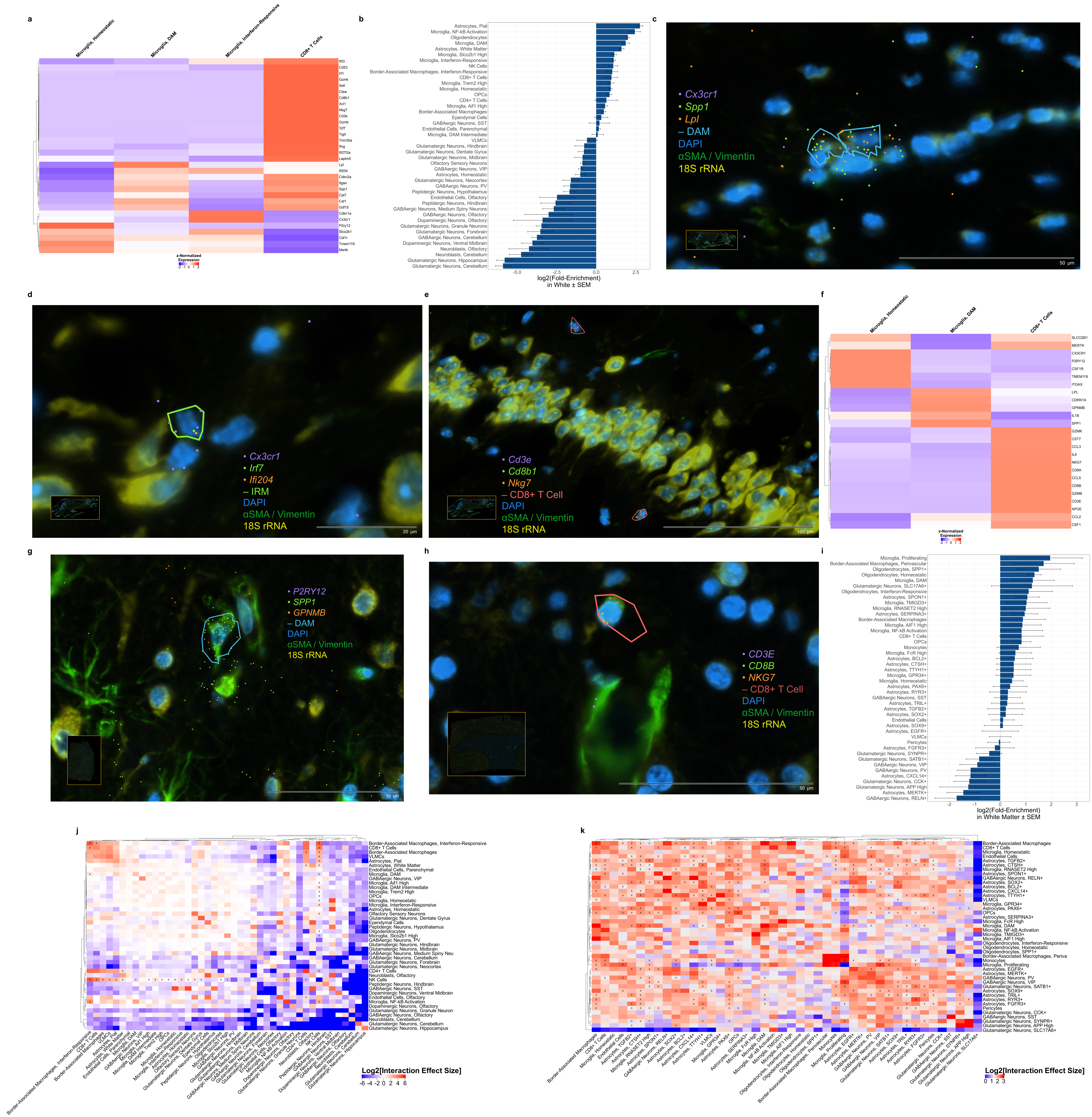

### Supplementary Figure S3

a

| IAV Gene | Total Counts in Dataset | Mean Counts per Cell | SD |
|---|---|---|---|
| HA | 0 | 0 | 0 |
| M1 | 0 | 0 | 0 |
| M2 | 0 | 0 | 0 |
| NA | 0 | 0 | 0 |
| NEP | 0 | 0 | 0 |
| NP | 0 | 0 | 0 |
| NS1 | 0 | 0 | 0 |
| PA | 0 | 0 | 0 |
| PA-X | 0 | 0 | 0 |
| PB1 | 0 | 0 | 0 |
| PB1-F2 | 0 | 0 | 0 |
| PB2 | 0 | 0 | 0 |

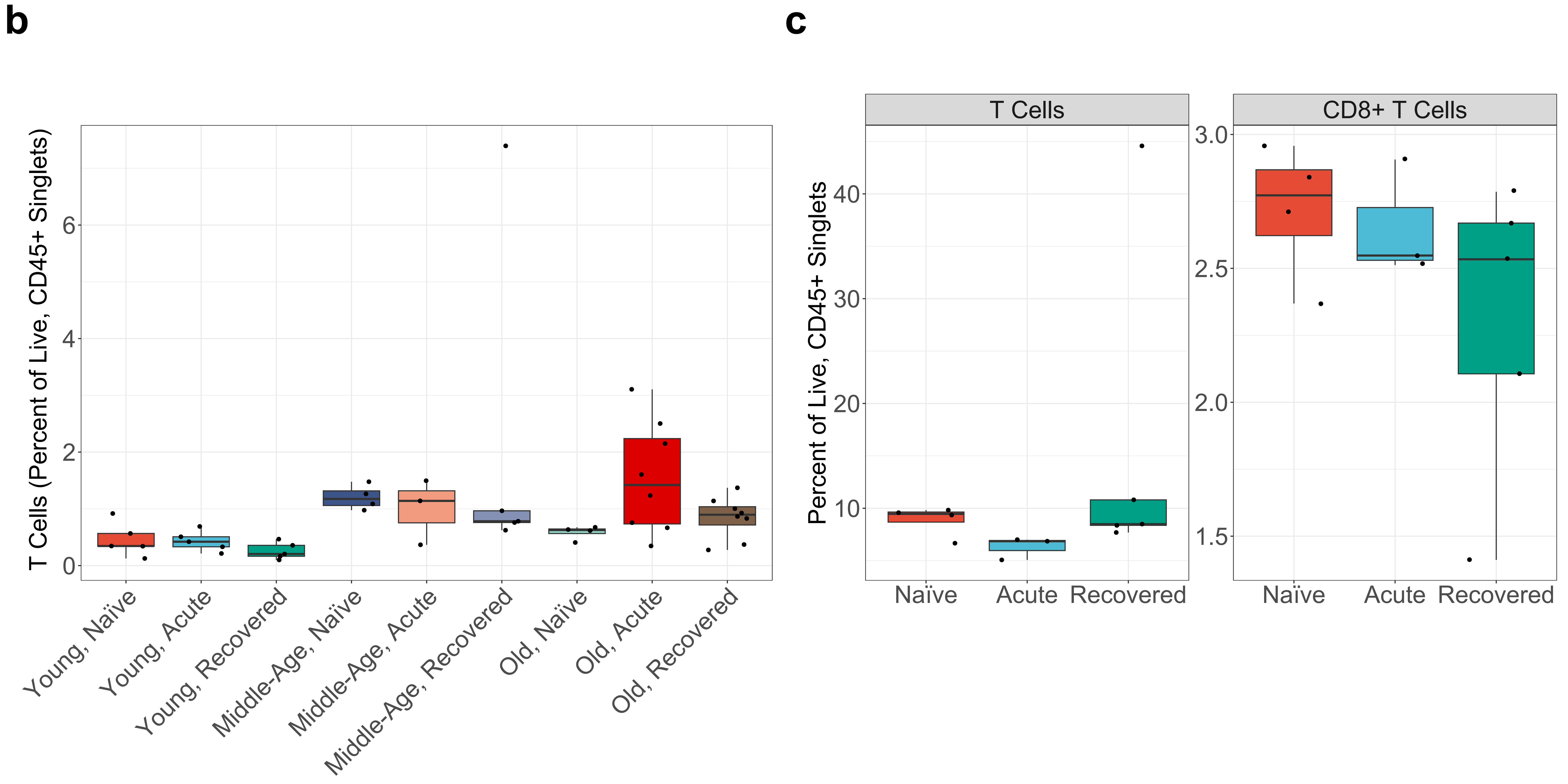


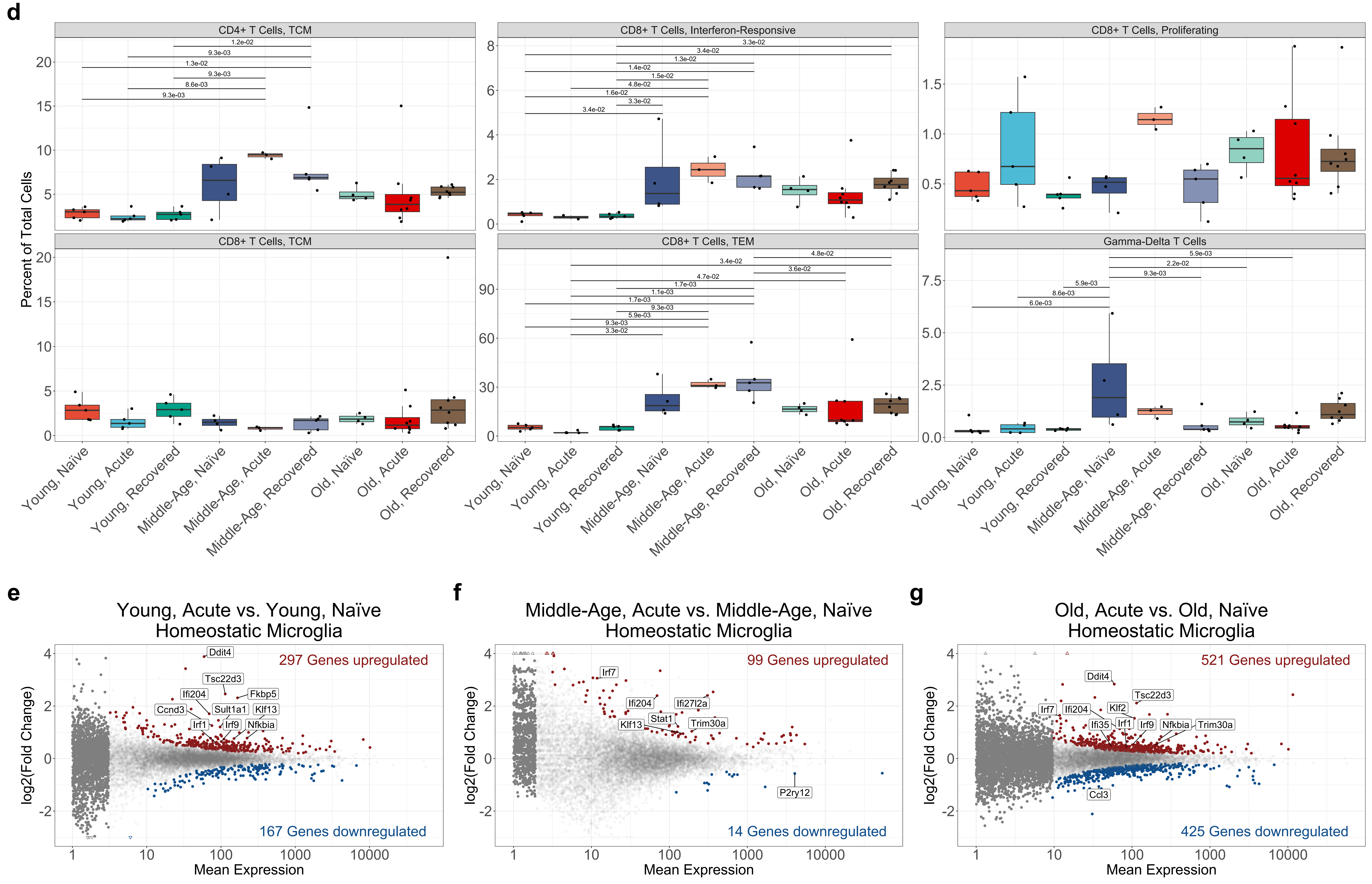

### Supplementary Figure S4

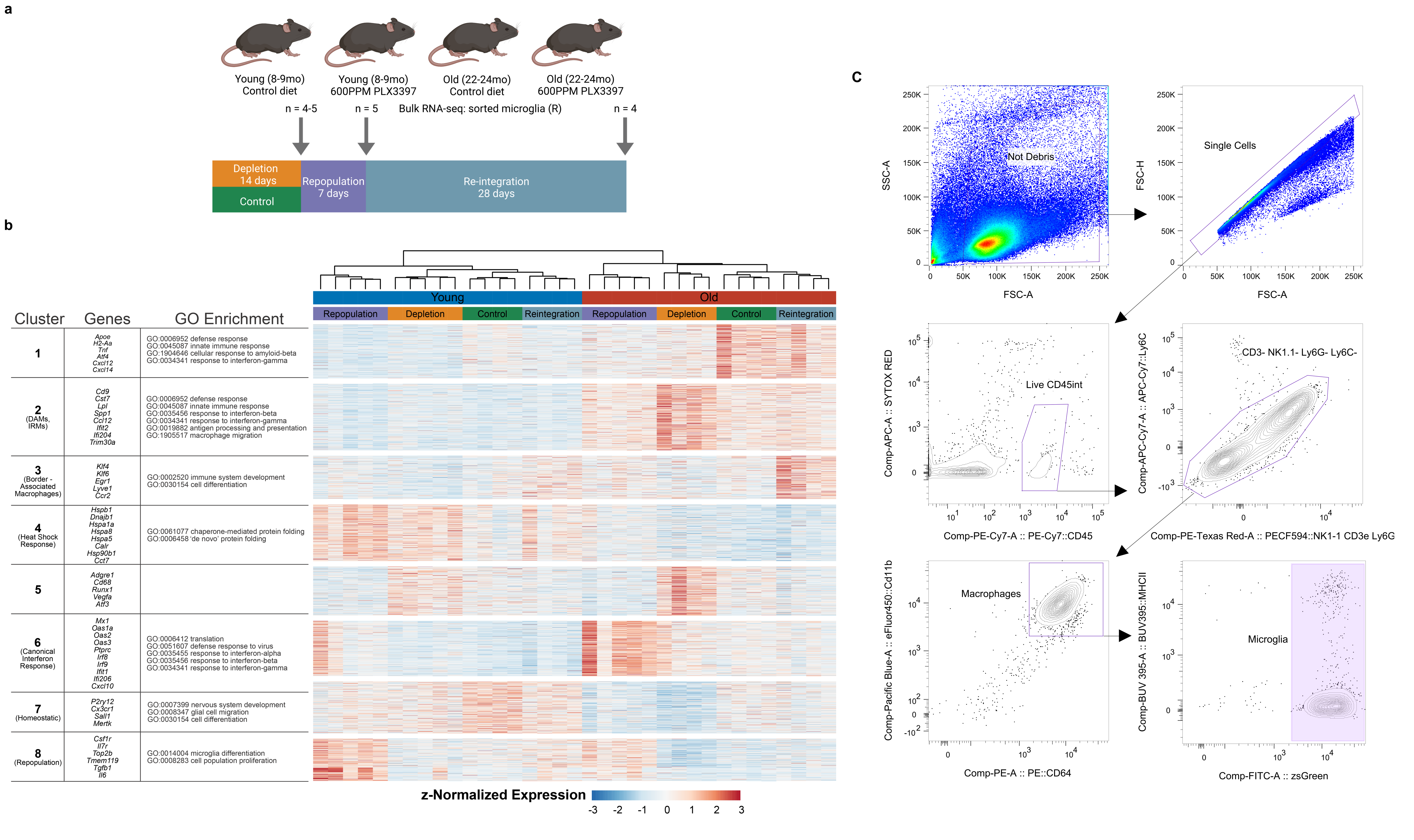
