## Supplementary Table S1 for "Influenza A infection accelerates disease-associated microglia formation during physiological aging"

Supplementary Table 4: Human brain aging cohort demographics

| Case Number | Clinical Diagnosis | Primary Neuropathologic Diagnoses | NIA–AA ADNC (ABC Score) | ApoE | PMI (Hrs) | Sex | Age at Death | Race/Ethnicity | Group |
|---|---|---|---|---|---|---|---|---|---|
| 1 | Amnestic dementia | Intermediate ADNC, LATE stage 2 | A3, B2, C2 | 3,3 | 8.5 | Female | 95 | Caucasian/non-hispanic | ADNC + LATE-NC, Old |
| 2 | SuperAger | Intermediate ADNC, LATE stage 2 | A3, B2, C3 | 3,3 | 20 | Female | 97 | Caucasian/non-hispanic | ADNC + LATE-NC, Old |
| 3 | SuperAger | Low ADNC | A1, B1, C1 | 3,3 | 6 | Male | 91 | Caucasian/non-hispanic | SuperAger, Old |
| 4 | Amnestic dementia | High ADNC, LATE stage 2 | A3, B3, C3 | 3,4 | 11 | Female | 75 | Caucasian/non-hispanic | ADNC + LATE-NC, Old |
| 5 | SuperAger | PART, LATE stage 2 | A0, B2, C0 | 3,3 | 16 | Male | 87 | Caucasian/non-hispanic | SuperAger, Old |
| 6 | SuperAger | PART | A0, B2, C0 | 3,3 | 9 | Female | 85 | Caucasian/non-hispanic | SuperAger, Old |
| 7 | Young control | None | None | Unavailable | Unavailable | Male | 28 | Unavailable | Normal Control, Young Adult |
| 8 | Normal control | Intermediate ADNC, LATE stage 1 | A3, B2, C1 | 3,4 | 21 | Male | 92 | Caucasian/non-hispanic | Normal Control, Old |
| 9 | Amnestic dementia | High ADNC | A3, B3, C3 | 3,3 | 16 | Male | 62 | Caucasian/non-hispanic | ADNC, Old |
| 10 | Amnestic dementia | High ADNC | A3, B3, C3 | Unavailable | 18 | Male | 66 | Caucasian/non-hispanic | ADNC, Old |
| 11 | Normal control | Low ADNC | A3, B1, C3 | 3,3 | 16 | Female | 87 | Caucasian/non-hispanic | Normal Control, Old |
